# Bottom-up reconstruction of minimal pyrenoids provides insights into the evolution and mechanisms of carbon concentration by EPYC1 proteins

**DOI:** 10.1101/2024.06.28.601168

**Authors:** A. M. Küffner, B. Pommerenke, L. Kley, J. Z. Y. Ng, S. Prinz, M. Tinzl-Zechner, P. Claus, N. Paczia, J. Zarzycki, G. K. A. Hochberg, T. J. Erb

**Affiliations:** Max Planck Institute for Terrestrial Microbiology, Karl-von-Frisch Str. 10, 35039 Marburg, Germany; Max Planck Institute of Biophysics, Central Electron Microscopy Facility, Max-von-Laue Str. 3, 60439 Frankfurt, Germany; SYNMIKRO Center for Synthetic Microbiology

## Abstract

Membraneless organelles play essential roles in cellular processes. In various photosynthetic organisms, they confer carbon concentrating mechanisms (CCMs). One example is the pyrenoid in *Chlamydomonas reinhardtii*, a liquid-phase separated organelle that localizes and improves carbon fixation via the intrinsically-disordered protein (IDP) EPYC1. Modern-day pyrenoids are complex structures, which makes it impossible to study the function of EPYC1, especially whether EPYC1 alone confers carbon concentration, and how EPYC1 could have initiated the evolution of pyrenoids. Here, we developed a bottom-up approach to study the function of EPYC1 and its sequence-function space across evolution. We demonstrate that modern-day EPYC1-sequences, but not other IDPs, induce liquid-phase separation of Rubisco into minimal pyrenoids with functional CCMs. Using ancestral sequence reconstruction, we trace the evolution of pyrenoids and demonstrate that selection acted on carboxylation rate, and in selected cases on specificity, providing unexpected perspectives on the design and function of natural and synthetic pyrenoids.

## Introduction

Membraneless organelles play key roles in organizing cellular processes in time and space across all domains of life^1–11^. In living cells, the role of these liquid-liquid phase separated structures, also called condensates, spans from RNA transportation, concentration buffering, to modulating metabolic networks at pivotal branching points^12–14^.

Notably, condensates are also involved in increasing the efficiency of photosynthesis. In various green algae, red algae, diatoms, and mosses, a condensate inside the chloroplast, called the pyrenoid functions as carbon concentrating mechanism (CCM)^8^. Pyrenoids are functionally conserved and evolutionary convergent structures that enrich inorganic carbon inside of photosynthetic cells^15,16^. They increase the local carbon dioxide concentration in close proximity to ribulose-1,5-bisphosphate carboxylase oxygenase (Rubisco), the CO_2_-fixing enzyme of the Calvin-Benson-Bassham cycle^17,18^. This local increase in CO_2_ promotes the carboxylation reaction of Rubisco and suppresses the unfruitful reaction of the enzyme with oxygen that causes the wasteful process of photorespiration^19^.

The best-studied case of condensate-based CCMs is the pyrenoid in the green algae *Chlamydomonas reinhardtii*, which is composed of several components. The inner part, where Rubisco is almost exclusively located, is called the pyrenoid matrix. This matrix is formed through co-condensation of the essential pyrenoid component 1 (EPYC1) protein, an intrinsically disordered protein (IDPs), with canonical form I Rubisco that consists of a small and a catalytically active large subunit (SSU and LSU)^20^. Matrix formation is essential for carbon concentration: EPYC1 mutants show no pyrenoid formation and cannot survive at ambient CO_2_ concentrations^8,21^. Beyond this functional core, CCM in *C. reinhardtii* contains additional elements: The matrix itself is further surrounded by starch sheets, which play a role in stabilizing the pyrenoid and are believed to prevent carbon leakage^9^. Additionally, thylakoid membrane tubules protrude into the pyrenoid that deliver inorganic carbon in form of HCO_3_^-^ for carbon fixation^22^. Within these tubules, carbonic anhydrase converts HCO_3_ ^-^ into CO_2_, which diffuses into the matrix, where it serves as substrate for Rubisco and is incorporated into ribulose-1,5,-bisphosphate to yield two molecules of 3-phosphoglycerate (3-PG).

However, it is still unclear, how pyrenoids function at a molecular level, and which minimal components would be required (or suffice) to reconstitute a functional CCM. This is even more interesting, when thinking about the convergent evolution of pyrenoids in green algae, red algae, diatoms and mosses, the diverse Rubisco forms that are targeted to the pyrenoid, as well as the structural organization and complexity of pyrenoids between these groups of organisms^23,24^

Recent work has delivered first insights into the molecular basis of pyrenoid formation in *C. reinhardtii*. *In vitro* studies showed that EPYC1 interacts with the SSU (but not the LSU) of form I Rubisco^10^ via short α-helical motifs to induce matrix formation ^25^. However, whether EPYC1-based condensation directly affects Rubisco catalysis, or whether the carbon concentrating effect of the pyrenoid is rather caused by secondary effects (i.e., substrate partitioning and/or delivery) is still unclear. In addition, what mechanisms could have driven formation of simple condensates, and what advantages such minimal condensates could have potentially provided, remains unanswered thus far.

Here, we took a synthetic biology and evolutionary biochemical approach to study the mechanisms and capabilities of different IDP-containing proteins on condensate formation *in vitro*. Using a reductionist approach, we demonstrate that EPYC1-homologs are sufficient to induce formation of minimal pyrenoids, when fused to Rubisco. We further show that pyrenoid formation is specific to EPYC1-homologs, while other (i.e., non-pyrenoid-related) IDPs do not promote phase-separation of Rubisco. Using ancestral sequence reconstruction, we traced pyrenoid formation and properties across the phylogenetic tree of EPYC1 to delineate evolutionary and ecological patterns in the EPYC1-sequence function space. Experimental characterization of resurrected minimal pyrenoids surprisingly show that evolution initially rather selected for carboxylation rate enhancement through EYPC1, and not selectivity. Finally, we demonstrate that EPYC1-based condensates enhance carboxylation without substantially altering the catalytic parameters or structure of Rubisco, and provide evidence that the basal CCM effect of EPYC1-based minimal pyrenoids is rooted in mass action kinetics. Together, these results demonstrate that EPYC1-sequences provide additional, carbon-concentrating functions, beyond phase separation, which can be used by natural and synthetic pyrenoids.

## Results

### EPYC1 causes condensation of form II Rubisco from *R. rubrum*

We first screened the capability of diverse IDPs for phase separation of Rubisco. To that end, we used the well-studied form II Rubisco from *R. rubrum* (CbbM) as target protein, and fused a library of IDPs^4,10,26–33^ to the protein’s C-terminus and N-terminus, respectively (figure 1 A, supplementary table 1). This library contained EPYC1 from *C. reinhardtii* (EPYC1_CR_), as well as 15 other IDPs that had shown condensate-formation as individual proteins, or when fused to other target enzymes before. Notably, when tested between 1 and 20 µM concentration, only the EPYC1 protein initiated functional and stable condensates with its non-natural target CbbM (supplementary figure 1, supplementary figure 2). All other IDPs showed no effect onto CbbM, except for a few that aggregated (supplementary figure 2 & 3). These experiments showed that specific sequences are required to generate pyrenoid like condensates *in vitro*, which is also indicated by the distinct sequence composition of EPYCs compared to other IDPs (supplementary figure 4 & 5).

**Figure 1.**
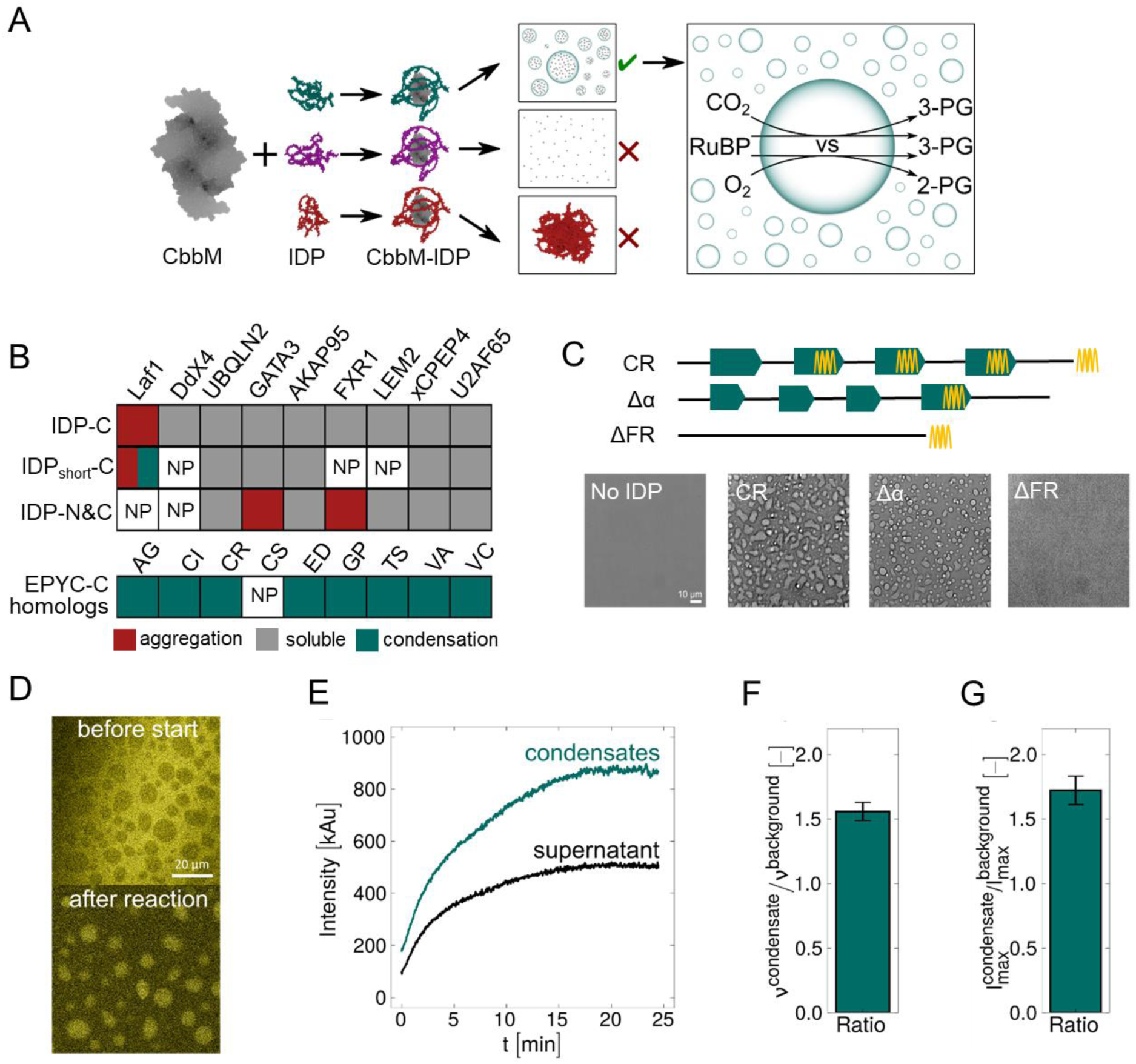
IDP-fusion screen to identify and characterize functional minimal pyrenoids *in vitro.* (A) General workflow. A library of IDPs^4,10,26–33^ was fused to CbbM (*R. rubrum* Rubisco). Each fusion protein was individually tested in a condensation screen for phase-separation, no effect (i.e., homogenous dispersal), or aggregation. Condensates were further subjected to carboxylation activity and specificity tests. (B) Overview of the fusion protein library and condensation screen results. Full (or shortened) IDPs were fused to the C-terminus (IDP-C, IDP_short_-C), or the N and C terminus (IDP-N&C) of CbbM. Red indicates aggregation, grey indicates homogeneously dispersed protein, white indicates not-producible (CbbM-Laf1N&C, CbbM-Ddx4N&C, CbbM-EPYC1_CS_), or not tested (CbbM-Ddx4_short_, CbbM-FXR1_short_, CbbM-LEM2_short_), green indicates condensate formation at concentrations of 1-20 μM. CbbM-Laf1_short_ undergoes liquid-liquid phase separation when mixed with 1 μM GFP and shows aggregation at all other conditions tested. (C) Architecture and behaviour of different EPYC1-IDP variants from *C. reinhardtii*. Teal arrows indicate repeat domains, yellow helices show SSU-recognition domains. CR = full length IDP, Δα = SSU-recognition domain deletion variant, ΔFR = repetition domains deletion variant. Light microscopy pictures show homogeneously dispersed CbbM and CbbM-EPYC1_CR_ΔFR, while CbbM-EPYC1_CR_ and CbbM-EPYC1_CR_Δα form condensates in 50 mM Tris (pH 7.5 & 8.0), 1 mM MgCl_2_, 5 mM NaHCO_3_, 0.1 mg/mL CA, and 0.3 mM RuBP. (D) Fluorescence microscopy pictures from the resorufin-coupled activity assay (supplementary figure 6) inside and outside of condensates before and after 25 minutes. (E) Fluorescence formation inside and outside of condensates. (F) Ratio of initial fluorescence formation rates between condensate inside and outside. (G) Ratio of maximum fluorescence levels between condensate inside and outside. Error bars represent standard errors from eight replicates.

We also tested truncated versions of EPYC1_CR_. Deletion of the full repeat domain of EPYC1_CR_ (EPYC1_CR_ΔFR) completely abolished condensate formation of CbbM, while truncation of the α-helical Rubisco small subunit (SSU) recognition motif (EPYC1_CR_Δα) did not change condensate-formation (figure 1 C). This showed that even without the SSU recognition motif, condensate-formation is in principle possible with simpler form II Rubiscos in our minimal system.

### EPYC1-based condensates are functionally active

We next tested, whether our synthetic condensates were functionally active and showed CCM properties. To assess the activity of CbbM within the condensates, we developed a coupled fluorescence assay (supplementary figure 6, material and methods). This assay catalyzes the oxidation of 3-phosphoglycerate (3-PG) to acetyl phosphate and hydrogen peroxide, which can be detected using horseradish peroxidase (HRP) and resorufin. We used confocal microscopy to monitor the fluorescence signal of CbbM-EPYC1_CR_Δα inside and outside of condensates over time (figure 1 D). In the absence of ribulose 1,5-bisphosphate (RuBP), the substrate of Rubisco, we observed only a weak background signal outside of the condensates. Upon addition of RuBP, we detected fluorescence formation inside and outside of the condensates. Notably, the rate of fluorescence formation was increased ∼1.6-fold inside the condensates and resulted in ∼1.5-fold increased total fluorescence levels. These findings showed that our condensates were enzymatically active and formed minimal “synthetic” pyrenoids that could (partially) emulate the function of natural pyrenoids (i.e., increased CO_2_ fixation).

To quantify the catalytic properties of EPYC1-condensed CbbM compared to homogeneous CbbM, we used radioisotope tracing with ^14^CO_2_. Because carbon concentrating mechanisms are most pronounced at sub-saturating substrate conditions (and our condensates additional showed limited stability at high substrate concentrations), we ran our assays at non-fully saturating substrate concentrations, that is 5 mM H^14^CO_3_^-^ ( ≈0.08 – 0.25 mM CO_2_) and 0.3 mM RuBP, and followed ^14^C-3-PG formation over time (figure 2 A & B). We determined carboxylation rates over the first 2 minutes, and quantified total ^14^C-3-PG levels at 90 min as proxy for carboxylation yield. In these experiments, phase-separated CbbM-EPYC1_CR_ showed carboxylation rates of about 4 CO_2_ s^-1^ per active site, and carboxylation yields of about 0.3 mM ^14^C-3-PG (≈50 % of RuBP). In strong contrast, the carboxylation rates of homogeneous (i.e., non-phase separated) CbbM were significantly (i.e., 4 to 5-fold) lower at 0.7-1 CO_2_ s^-1^ per active site. Additionally, homogeneous CbbM formed only 0.2 mM ^14^C-3-PG product (≈33 % of RuBP). Overall, this data indicated that CbbM showed lower specific activities and catalyzed a substantial fraction of oxygenation, when not phase separated into a pyrenoid, which we also confirmed experimentally subsequently (see below).

**Figure 2.**
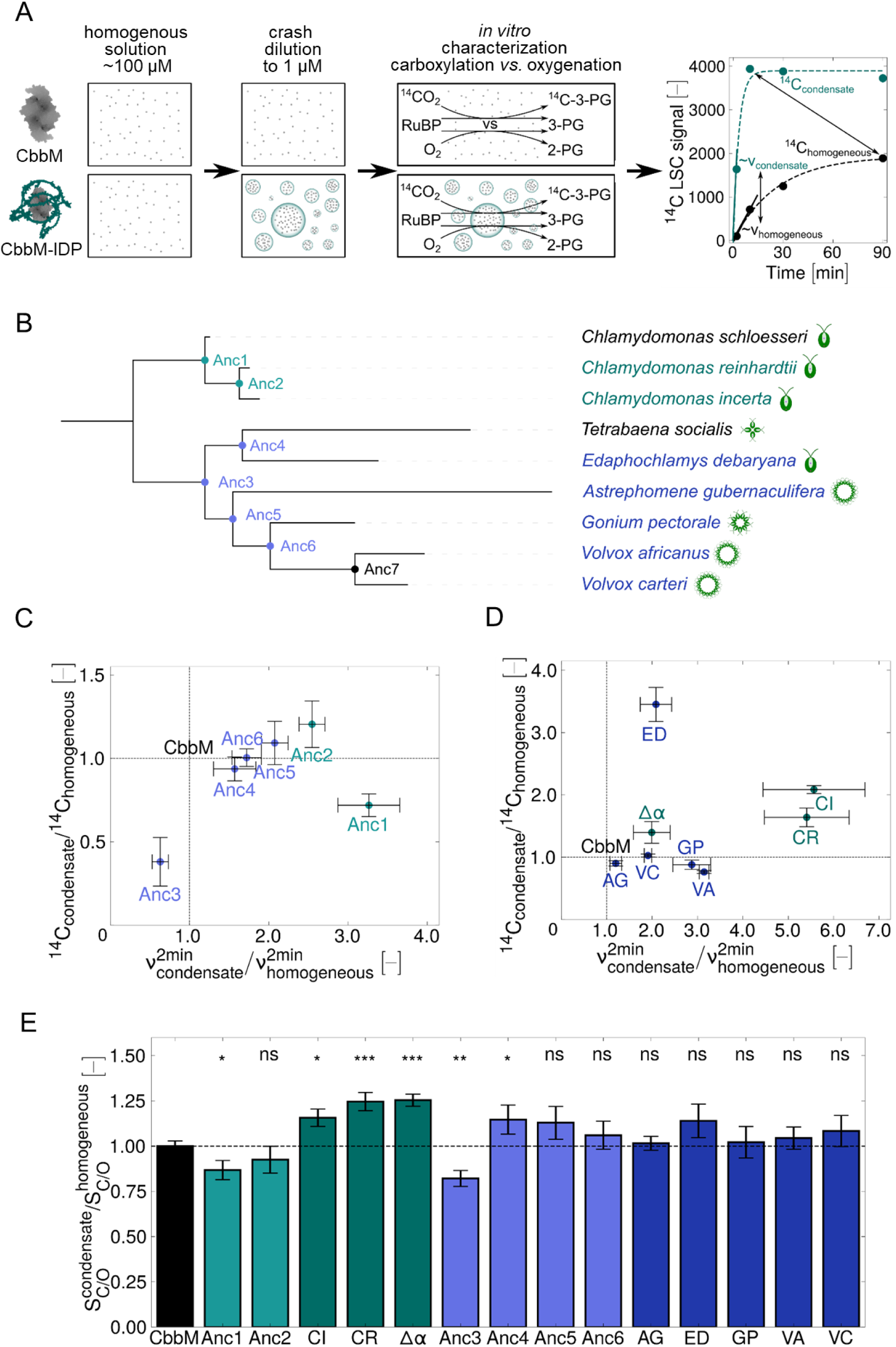
Characterization of EPYC homologs across the phylogenetic and evolutionary tree. (A) General workflow. CbbM and CbbM-IDP fusion proteins were prepared at ∼100 µM concentrations (yielding homogenously dispersed protein stocks). Crash dilution to 1 µM final concentration was used to induce phase separation of the CbbM-IDP fusions. Carboxylation assays were performed side-by-side at 25°C using ^14^C-CO_2_, yielding time curve data of homogenous CbbM (black lines) and phase separated CbbM-IDP (green lines). Arrows indicate the computation of initial carboxylation rate ratios and maximum ^14^C incorporated ratios. (B) Phylogeny of EPYC1_CR_ and its nine known homologs as inferred by maximum likelihood. The tree was rooted at the oldest node following the species relationships of the *Chlamydomonales* order^34^. Teal and light purple show cloned, produced and stable EPYC1 ancestors; green and blue indicate cloned, produced and stable extant EPYC1 homologs. EPYC1_CS_ could not be subcloned, EPYC1_TS_ forms condensates in buffer that dissolve under reaction conditions, Anc7 showed little protein production and no phase separation. (C) Maximum ^14^C measured ratios versus initial rate ratios for the produced fusion proteins of CbbM and EPYC1 ancestors. Each ratio was computed from a ^14^C time curve of CbbM-IDP in condensate-forming conditions compared to CbbM only. Ancestor fusion proteins from the *Chlamydomonas* family are shown in teal, and ancestor fusion proteins from the *Volvox* family in light purple. The crossed lines delineate the behaviour of homogeneous CbbM. Reactions were conducted in non-saturating conditions at pH 7.5-8.0, 5 mM NaHCO_3_ or NH_4_CO_3_, 0.1 mg/mL CA, 0.3 mM RuBP, 25 °C, 300 rpm, and at atmospheric conditions. For each ratio, at least three independent replicates for homogeneous CbbM, and phase-separated CbbM-IDP were used. The variance of homogeneous CbbM was estimated from all CbbM measurements. For EPYC_CR_ two datasets were combined with two and three replicates, respectively. For EPYC_ED_ two data sets of each three replicates were combined. Variance was computed using empirical error propagation. Error bars indicate standard errors. (D) Maximum ^14^C measured ratios versus initial rate ratios for the produced fusion proteins of CbbM and EPYC homologues. Data sets were treated the same as in D. Error bars indicate standard errors. (E) Ratios of specificity between homogeneous CbbM and phase-separated CbbM-IDP measured in 4991 ppm CO_2_ in O_2_. In each specificity assay CbbM was measured alongside the fusion protein to control for daily variations. Errors were propagated using empirical error propagation. Error bars indicate standard errors from at least three biological replicates. P-values were computed using a two-sided t-test assuming independent variances (p-value<0.05=*, <0.01=**, <0.001=***).

### Condensate-forming properties are conserved across EPYC1 phylogeny

In the following, we sought to investigate the potential of homologs of EPYC1 to form minimal pyrenoids. So far, nine EPYC1 homologs are known from green algae, all from the order of *Chlamydomonadales*. Of these nine sequences, we could successfully produce eight as fusion proteins with CbbM. All eight homologs formed condensates (supplementary figure 2), showing that the basic condensate-forming property is conserved across these homologs, despite a sequence conservation as low as 57% between the different variants (supplementary table 3 & 4).

To study the evolution of these sequences, we created a phylogeny of the nine EPYC1-containing species in an unrooted tree (supplementary figure 7), which we subsequently rooted using the known species relationships of *Chlamydomonadales*^34^ (figure 2 B). In our phylogeny two main families diverge within the *Chlamydomonadales*: the *Volvox* family and the *Chlamydomonas* family. Within the *Chlamydomonas* branch, all species are strictly unicellular fresh-water organisms, whereas almost all species of the *Volvox* branch inhabit fresh and salt water and form multicellular spheroidal colonies, with exception of the unicellular species *Edaphochlamys debaryana*^35–38^. We used ancestral sequence reconstruction to infer ancestral EPYC1 sequences at internal nodes along the phylogenetic tree (Anc1-7) and expressed them as C-terminal fusions to CbbM. Except for Anc7, which exhibited minimal protein production and condensate formation, all ancestral sequences resulted in stable proteins that induced phase separation of CbbM. This data suggested that the condensate-forming property was already prevalent in all ancestral EPYC1 sequences, including the last common ancestors of the *Chlamydomonas* (Anc1), as well as the *Volvox* family (Anc3), respectively.

### Ancestral EPYC1s enhance catalysis in condensates

In the following, we tested the extant and ancestral sequences in respect to carboxylation rate enhancement and maximum amount of 3-PG produced, using the radioactive label test described above. Notably, all ancestors (with exception of Anc3) exhibited 2- to 3-fold increased rates compared to homogeneous CbbM at sub-saturating conditions, suggesting that rate enhancement was an ancestral trait (figure 2 D). With respect to maximum 3-PG formation (as proxy for specificity), the picture was more complex. Compared to homogeneous CbbM, the last common ancestors (Anc1 and Anc3) showed even decreased 3-PG production, while 3-PG production was not significantly different in all other ancestors (and only slightly enhanced in Anc2), indicating that ancestral EPYC1s did initially not confer specificity advantages.

When expanding to extant homologs of EPYC1 (figure 2 D), this allowed us to infer general trends for rate enhancement along the evolutionary trajectories of the *Chlamydomonas* and the *Volvox* family, respectively. In the *Chlamydomonas* branch, Anc1 and Anc2 had already enhanced carboxylation rate 2-3-fold in condensates. This effect was even increased (5- to 6-fold) for the extant EPYC1-homologs of *C. reinhardtii* and *C. incerta* (figure 2 D), indicating that rate enhancement was selected for by evolution. Within the *Volvox* branch, we observed a similar pattern. While Anc3 showed decreased carboxylation rates, we observed 2- to 3-fold enhanced rates for ancestors Anc4-6 in condensates, and all extant homologs of the *Volvox* family (except for *Astrephomene gubernaculifera*).

With respect to the maximum amount of 3-PG produced, of the ancestors in the *Chlamydomonas* branch, Anc2 showed an improved ratio of ∼1.6-fold and this ratio was comparable to all extant *Chlamydomonas* homologs. In the *Volvox* branch, only the extant homolog from *E. gabarayana* showed ∼3-fold increased maximum production of 3-PG. This is interesting, as *E. gabarayana* is the only unicellular species of all EPYC1-encoding organisms within the *Volvox* family, eventually pointing to an evolutionary adaptation of this EPYC1-homolog that reminds that of EPYC1 homologs in the unicellular *Chlamydomonas* family, which also show increased specificity^35,38^.

To quantify specificity more precisely, we used defined gas mixtures and determined the formation of (3-phospho) glycerate and (2-phospho) glycolate, respectively (figure 2 E). These data showed in general comparable trends to those of the maximum 3-PG formation assay, but overall lower effects on specificity. This was probably caused by the differences between the two assays, specifically, the supply of inorganic carbon, which is provided in the form of bicarbonate in case of the maximum 3-PG formation assay, and as CO_2_/O_2_ gas mixture in case of the specificity assay.

Altogether, our data showed that ancestral EPYC1-homologs possessed phase-separating capabilities down to the last common ancestors, and consistently enhanced carboxylation rates in condensates. In contrast, specificity-conferring properties that are directly encoded in the EPYC1-sequence seemed to have evolved only within some algae lineages, likely correlating with specific lifestyles (i.e., unicellularity).

### Substrate partitioning and mass action effects explain CCM

Having demonstrated that EPYC1 caused functional CCM *in vitro* (i.e., increased carboxylation rate and specificity), we next thought to explain the underlying mechanism. There are two main models that are currently considered. According to one model, condensates change the catalytic output of a reaction mixture by directly affecting the kinetic properties of enzymes^39^. The different chemical (e.g. a less polar) environment within the condensate is assumed to cause conformational or allosteric effects and change the preferred reaction pathway of a given enzyme^39^, similar to the influence of organic solvents on the lid behavior and catalysis of esterases/lipases^40^. In the alternative model, physical-chemical effects, such as substrate partitioning are at work (figures 2 B & C, 3 A & D). In this case, the different solvation of substrates causes the concentration of certain reactants inside of condensates to vary, resulting in multiphase reaction systems with mass action effects that affect catalytic rate and specificity.

**Figure 3.**
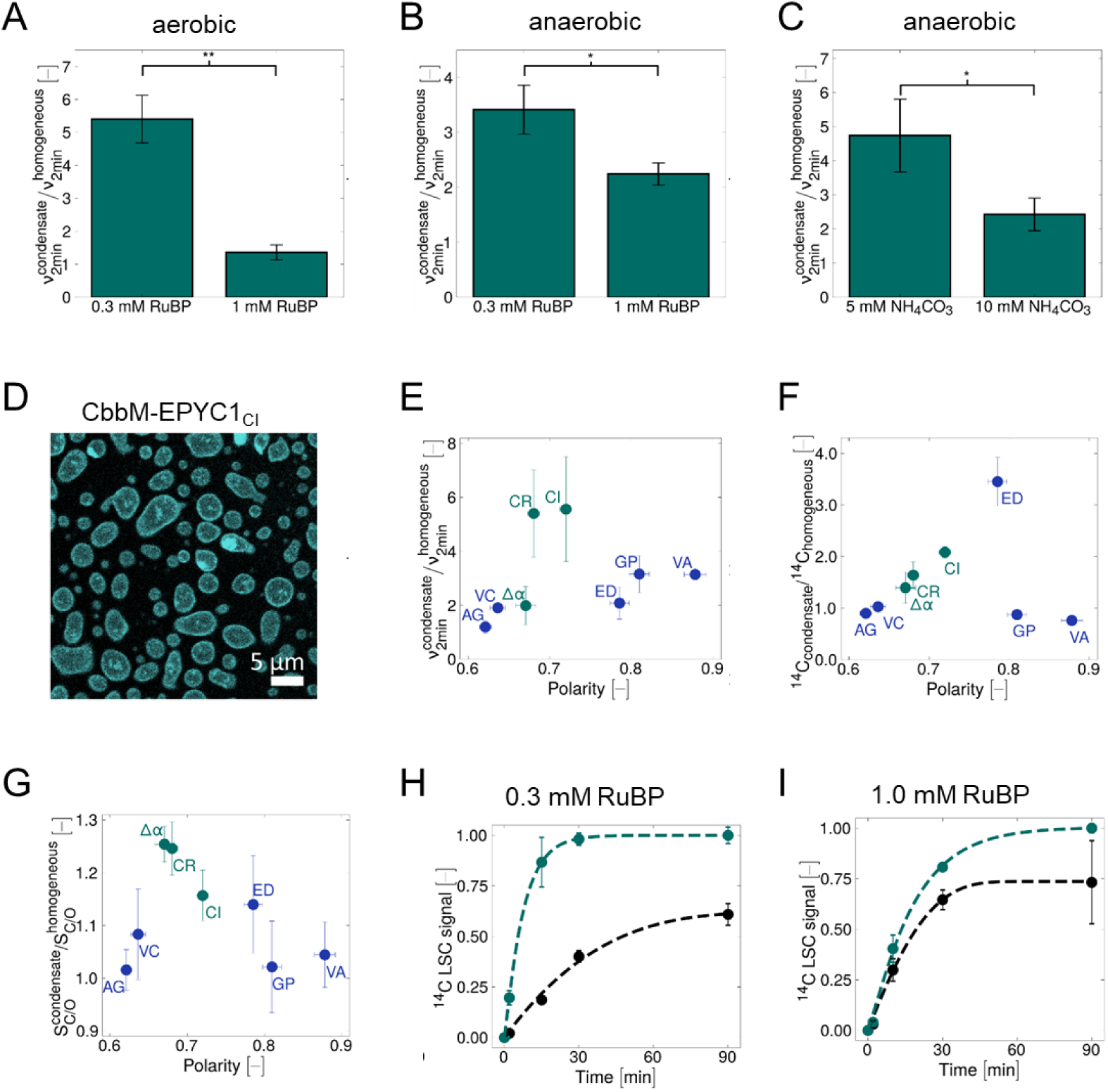
CbbM-IDP kinetics and material property data of condensates. (A, B) Ratio of initial carboxylation rates between CbbM and CbbM-EPYC1_CR_ at 0.3 and 1 mM RuBP at aerobic (A) or anaerobic (B) conditions. (C) Ratio of initial carboxylation rates between CbbM and CbbM-EPYC1_CR_ at 5 mM and 10 mM CO_3_^2-^ at anaerobic conditions. Microscopy analysis showed no morphological difference between the different conditions used in A-C (supplementary figure 8). Ratios were determined from at least three independent replicates. P-values were computed using a two-sided t-test assuming independent variances (p-value<0.05=*, <0.01=**, <0.001=***). (D) Exemplary confocal microscopy image of condensates formed by CbbM-EPYC1_CI_ with partitioned solvatochromatic dye PRODAN (∼1 mM, pH 7.5, 50 mM Tris) to quantify polarity of condensates^42^. (E) Ratios of initial carboxylation rates of different EPYC1-condensates plotted against condensate polarity. (E) Maximum ^14^C incorporated ratios from figure 2 C & D plotted against condensate polarity. (G) Specificity ratios from figure 2 F plotted against condensate polarity. Polarity was measured from more than 10 condensates each in at least 3 biological replicates. (H) Data fitting of mass action-mass transfer models to the time curve data in aerobic conditions at 0.3 mM RuBP from condensates formed by CbbM-EPYC1_CR_ (teal) and from homogeneous CbbM (black). (I) Data fitting as in H to data obtained with 1 mM RuBP.

To distinguish between these two hypotheses, we utilized the fact that mass action kinetic effects typically decrease with increasing substrate concentrations^41^. This reduction is not due to increased specific catalytic activity (v_max_), but increased active site saturation at increasing concentrations, and thus convergence of catalytic rates (k_cat_E limit). Indeed, increasing RuBP concentrations (under aerobic and anaerobic conditions; figure 3 A & B), as well as increasing CO_2_ concentration (under anaerobic conditions, figure 3 C) decreased the differences in rate enhancement between phase-separated and homogenous CbbM, suggesting a mass action-based CCM in our minimal pyrenoids.

To independently test both hypotheses, we used a model-based approach, in which we fitted mass action kinetic models based on generalized Michaelis-Menten kinetics^41^ to our experimental data at 0.3 and 1.0 mM RuBP (figure 3 H & I). To test for the substrate partitioning hypothesis, we varied the partitioning of RuBP, CO_2_, and O_2_ into the condensates, while keeping the kinetic parameters of CbbM constant for both phase-separated and homogeneous conditions. With these assumptions, our experimental data was modelled best by a ratio of CO_2_/O_2_ partitioning between 3.6 to 8.5 (at 0.3 and 1 mM RuBP, respectively, supplementary table 2), which is well in line with previously reported partitioning coefficients^42,43^.

We also tested the alternative hypothesis that condensate formation directly affects the kinetic properties of CbbM by varying K_m_(CO_2_) and K_m_(O_2_) for phase-separated conditions. Varying those parameters, the experimental data could be only explained by a 6-fold (1.5-fold) decrease in K_m_(CO_2_) and a 14-fold (2.6-fold) increase in K_m_(O_2_) at 0.3 (1 mM) RuBP inside the condensates. However, this would result in a dramatic change in CbbM specificity within the condensates between one and three orders of magnitude at 1 mM and 0.3 mM RuBP, respectively. Such direct change of enzymatic parameters through condensate formation is unreported, and much less likely compared to the substrate partitioning hypothesis, which we further supported by structural analysis of Rubisco in condensates (see below).

To further elucidate the physical-chemical mechanisms of CCM, we investigated the material properties of our condensates. We used a previously developed confocal microscopy-based polarity assay^42^, to measure the polarity inside the different minimal pyrenoids (figure 3 D, supplementary figure 9). This data indicated a slight optimum for rate enhancements around a polarity of 0.7, and a more pronounced optimum for 3-PG formation and specificity, respectively between 0.65 and 0.8 (figure 3 E & G). This optimum reminds of the Sabatier^44^ principle in heterogeneous catalysis, where adsorption and desorption of substrates and products on the catalyst are determining the catalyst performance in form of a sharp, volcano-plot like behaviour.

### Structure analysis shows CbbM dimer formation in pyrenoids

Finally, we further investigated whether condensate formation would have any structural or allosteric effects onto CbbM. To that end, we first solved the crystal structure of homogenous CbbM (without EPYC1) in presence of the substrate analog 2-carboxyarabinitol-1,5-bisphophate (CABP) at a resolution of 2.5 Å (figure 4 A). We then aimed at obtaining the structure of CbbM from condensates through single particle reconstruction (SRP) from cryogenic transmission electron microscopy (cryoTEM) images of phase-separated CbbM in the presence of CABP. To that end, we used the CbbM-EPYC1_CR_Δα construct that lacks most of the SSU-binding domains (figure 1), but retained catalytic properties of full-length EPYC1_CR_ (figure 2, supplementary figure 12).

**Figure 4.**
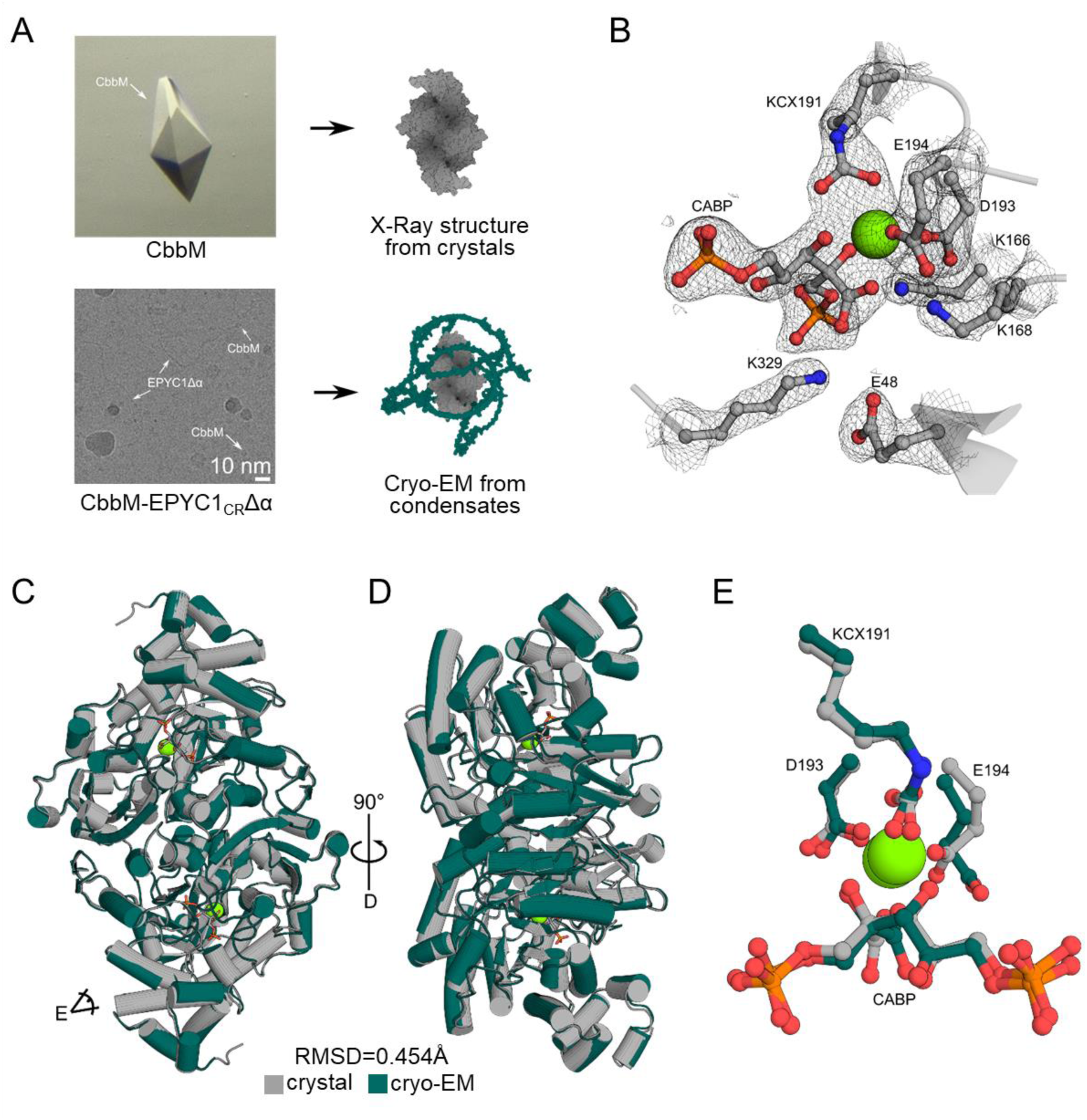
Structural analysis of crystallized and phase-separated CbbM. (A) General workflow for structural comparison of crystallized CbbM and phase separated CbbM-EPYC1_CR_Δα. CbbM was purified and crystallized via vapour diffusion in an open and closed conformer upon binding of CABP. CryoEM images of phase-separated were recorded at 0.655 Å/pixel resolution in ∼50 mM final NaCl (condensate condition) at the edges of condensates. (B) Active site of crystallized CbbM in the crystal at 2.5 Å. (C) Crystal structure of CbbM (grey) superimposed with the cryoEM structure of CbbM-EPYC1_CR_Δα (green). (D) Superimposed model (C) rotated by 90° (E) Zoom on the overlay of the inhibitor CABP, the carbamylated lysine K191 (KCX) and the Mg^2+^ between the crystal structure (grey) and the cryo-EM structure (green) of CbbM.

We formed condensates on the carbon grid and collected TEM images at the edges of condensates, where density and thickness of the protein layer still allowed for single particle reconstruction. This approach allowed us to solve the structure of the complete, phase-condensed CbbM domain at a resolution of 3.4 Å. Notably, we exclusively detected dimeric particles, but no higher-order oligomers. We also did not detect any additional density, suggesting allosteric binding of parts of the EPYC1_CR_Δα to the CbbM itself. This is fully in line with mass photometer data of homogenous CbbM and homogenous CbbM-EPYC1_CR_Δα (in non-condensate forming buffer) that both show a dimeric state of the enzyme (supplementary figure 13), indicating that the CbbM domain also formed a native complex in the condensate. Moreover, the CbbM structure from the condensates overlaps perfectly with the crystal structure of CbbM (figure 4 C-E, RMSD value of 0.494 Å over 6051 atoms), with almost perfect alignment of their active sites, indicating only minimal structural changes between condensed and homogenous CbbM. While a slight twist in E194 and bound CABP in the active site (figure 4 E) was observed, this was likely caused by the lower resolution of the condensate structure and the best-fit procedure, and/or radiation damage from cryoEM.

Besides the (dimeric) CbbM particles, we also observed long, winding rods in the TEM images that we suspected to represent the EPYC1_CR_Δα-proteins. Although we were unable to reconstruct those rod-like shapes together with the CbbM domain at high resolution, we could obtain some 2D classes of these structures. These structures resembled one to several EPYC1_CR_Δα-proteins in a bundled shape that we were able to reconstruct to resolutions of approximately 12 Å (supplementary figure 11). Together, this data suggested that the EPYC1_CR_Δα-proteins bundled CbbM into the condensate, where the CbbM domain adopted its native dimeric state. In summary, our structural analysis supported the hypothesis that the CbbM domain was not allosterically modified in the pyrenoid, and that the observed CCM of EPYC1 was mainly caused by mass action kinetics.

## Discussion

Understanding the molecular mechanisms of CCM in pyrenoids is a fundamental challenge, which has been addressed mainly by *in vivo* studies in a top-down fashion. Here, we used a reductionist, bottom-up approach to study pyrenoid formation *in vitro* from a mechanistic, evolutionary and structural point of view. Using a simple proteobacterial form II Rubisco (CbbM), we show that fusion with modern-day EPYC1-sequences is sufficient to induce phase separation of CbbM into minimal pyrenoids, and that (some of) these condensates possess carbon concentrating properties. This suggests that EPYC1-proteins already encode basic properties for enhanced carboxylation rates – and in certain cases – even more specific CO_2_-fixation. Yet, we note that our observation does not imply that additional factors and/or proteins could not further enhance CCM in natural pyrenoids. As a matter of fact, for optimal function of pyrenoids, a more complex and dynamic organization might be required, especially *in vivo*.

Although modern-day pyrenoids are more complex structures, our results provide evidence that for the initial evolution of pyrenoids, a single phase-separating protein could already provide a benefit and was likely sufficient. This allows to draw an evolutionary path from minimal pyrenoids to more complex structures, which are prevalent in green algae today. Considering the relatively high levels of CO_2_ (and low levels of O_2_) during early Earth history it is unlikely that selection initially acted on enhanced specificity in EPYC1-sequences. This is in line with our data on ancestral (and extant) EPYC1-homologs that consistently show carboxylation rate enhancements, indicating that this feature of EPYC1 was selected for throughout evolution. The evolution of enhanced specificity, which is more relevant today, appears to be a more recent phenomenon, likely driven by declining environmental CO_2_ concentrations (and increasing O_2_ concentrations), and apparently particularly in *Chlamydomonales* representatives with unicellular lifestyle.

Through structural studies, theoretical and material analysis, we propose that the CCM of EPYC1 in our minimal pyrenoids is mainly based on substrate partitioning and mass action kinetic effects, rather than direct effects of the pyrenoid matrix onto the structure and/or catalytic properties of CbbM. This highlights the significance of physical-chemical principles in the basic function of carbon concentration and opens the possibility for future applications: Our approach to use simple phase-separating IDPs to establish minimal carbon concentrating structures successfully conclude recent protein design efforts with similar goals^45^, and provide an alternative strategy to complex pyrenoid reconstruction efforts and/or transplantation approaches aimed at enhancing carbon fixation.

## Acknowledgements

The authors thank the Central Electron Microscopy Facility at the Max Planck Institute of Biophysics for expertise and access to their instruments. We acknowledge support from the EMBL Hamburg at the PETRA III storage ring (DESY, Hamburg, Germany). A.M.K gratefully acknowledges the help of Luca Schulz in the radioisotope laboratory, and Franziska Sendker for the introduction to the mass photometer.

## Funding

A.M.K. is grateful for funding provided by EMBO (ALTF 684-2022) and MSCA (Project 101106795 — ECOFix). A.M.K., B.P, L.K, J.Z.Y.N., S.P., M.T., P.C., N.P, J.Z., G.K.A.H., and T.J.E. are grateful for the generous support from the Max Planck Society.

### Author contributions

A.M.K conceived the project, analyzed data, and planned experiments. A.M.K, B.P, L.K performed molecular cloning and protein purification. A.M.K performed all activity, light and confocal microscopy experiments, mass photometer measurements and model simulations. J.Z.Y.N., G.K.A.H., and A.M.K. performed the ancestral sequence reconstruction. M.T. conceived and optimized the fluorescence Rubisco carboxylation assay. A.M.K., P.C., and N.P. optimized, prepared and measured the LC-MS/MS methods and samples. S.P. prepared and froze cryoEM grids and acquired cryoEM data. A.M.K. processed cryoEM data. A.M.K. and J.Z. crystallized CbbM, collected and processed x-ray diffraction data. J.Z. solved and refined crystal and cryoEM structures. T.J.E. supervised the project. A.M.K. and T.J.E. wrote the manuscript with contributions and comments from all authors.

## Materials & Methods

### Reagents and materials

Unless otherwise stated, chemicals were obtained from Sigma-Aldrich and Carl Roth in the highest purity and quality. NaH^14^CO_3_ and K^14^CN were obtained from Hartmann Analytics (Germany) and PerkinElmer (MA). Reagents for cloning were obtained from New England Biolabs, Macherey-Nagel, Qiagen, and Thermo Fisher Scientific. For first tests, commercial D-Ribulose 1,5-bisphosphate sodium salt hydrate (>90% purity, R0878, Sigma-Aldrich) was used. For detailed characterizations, enzymatically synthetized RuBP following published protocols^46^ was employed. For the determination of specificity constants, [1-3H] RuBP was synthetized enzymatically from D-[2-^3^H] glucose (PerkinElmer, MA) following published protocols^47^. The Rubisco inhibitor 2-carboxyarabinitol-1,5-bisphosphosphate (CABP) was previously synthetized in two batches using either cold or [^14^C]-labeled KCN using published protocols^46^. The concentration of the cold CABP batch was previously determined by inhibiting a Rubisco stock containing a known active site concentration (as quantified by [^14^C]-CABP) with varying concentrations of cold CABP and used with the stated concentration. Unless otherwise stated all data was analyzed and processed using python 3.10 with packages matplotlib^48^, pandas^49^, scipy^50^, numpy^51^, openCV^52^,scikits-odes^53–55^, and statsmodels^56^.

### Phylogenetics and ancestral sequence reconstruction

To infer the evolutionary relationship of EPYC1 and homologs, amino acid sequences were gathered by online BLASTP on 06 September 2022 using the amino acid sequence of EPYC1 from *C. reinhardtii* (XP_001690584.2) as the initial query. Sequences were aligned using MUSCLE (v.3.8.31)^57^ followed by tree inference by IQ-TREE (v.2.2.0.3)^58^ and a best fit model of evolution as determined by Modelfinder^59^, with gamma-distributed among site rate variation and fixed base frequencies. Two models of evolution were identified by IQ-TREE and tested. First Mitochondrial Inverterbrate (mtInv)^60^ in case of considering all amino acid evolution models and secondly, Jones-Taylor-Thornton (JTT)^61^ in case of restricting to the most commonly chosen amino acid models of LG^62^, JTT, and WAG^63^. Both approaches led to the same inferred sequences with respect to the natural variation (supplementary tables 3 & 4).

Ancestral sequences and posterior probabilities (PP) of ancestral states were reconstructed at internal nodes as implemented in IQ-TREE, using the same evolutionary model as per tree inference. Gap assignment of ancestral sequences was determined using Fitch parsimony with PastML (v.1.9.34)^64^ yielding our final tree. Ancestral sequences contain states with the highest PP at all sites selected.

The resulting evolutionary tree was rooted using timetree.org^34^ to the oldest node. Estimates of atmospheric gas concentrations were directly adapted from the website.

### Protein production and purification

EPYC like protein coding DNA was codon optimised and produced by Twist or Azenta Biosciences. Constructs were cloned to the C-terminus or N & C-terminus of the *R. Rubrum* CbbM coding region in a His-bdSUMO plasmid acquired from Addgene^65^ using OneShot^TM^ Top10 cells (Invitrogen). The constructs were expressed in OneShot^TM^BL21-AI^TM^ (Invitrogen) cells in TB media. An ON seed culture was used to inoculate 6L of production culture. Cells were harvested by centrifugation at 5000 g in a Beckman Coulter Avanti JXN-26 centrifuge equipped with a JLA-8.100 rotor. The cells were resuspended in 50 mM Tris, 10 mM MgCl_2_, 500 mM NaCl, 2 mM 2-BME, 10 mM Imidazole at pH 8.5. Cells were broken using a microfluidizer at 18000 psi with 4 cycles. Subsequently, the cell debris was sedimented using ultracentrifugation at 100000xg (Sorvall^TM^ WX+, Thermo Scientific with A-27 fixed angle rotor 6 x 50mL). The supernatant was then filtered using 0.2 nm syringe cap filters. The filtrate was subsequently applied to a 50 or 150 mL superloop (Cytiva, MA) and from there on 2 stacked 5 mL HisTrap columns (Cytiva, MA). The protein on the column was washed with 50 mM imidazole, 50 mM Tris, 500 mM NaCl, 2 mM 2-BME, pH 8.5 until the UV 280nm signal was stable. Subsequently, 5 mL of 2 mg/mL bdSNEP1 in 50 mM imidazole, 50 mM Tris, 500 mM NaCl, 2 mM 2-BME, pH 7.5 supplemented with 2mM DTT was applied to the columns and incubated for 5 hours at 20°C. Then the cleaved protein was eluted using 10 mL of 50 mM imidazole, 50 mM Tris, 500 mM NaCl, 2 mM 2-BME , pH 8.5 and captured in the superloop. Subsequently, the protein was applied to a 16/600 200 pg Superdex SEC-column in 50 mM Tris, 500 mM NaCl, 10 mM, MgCl_2_, pH 8.5 with 10 % v/v glycerol, isocratically eluted, and 2mL fractions were collected. The fractions were analysed for protein content and impurities using SDS-PAGE and then concentrated to ∼100 μM using 10kDa or 30kDa MWCO concentrators (Amicon® Ultra Filters, Merck). The protein was aliquoted into 20 μL volumes, frozen in liquid nitrogen and stored at -80°C until further use. For cryoEM structure elucidation within condensates, the purest SEC fraction (w/o glycerol) of CbbM-EPYC1Δα was used. For crystallisation, the CbbM fraction with the highest purity from SEC using 20 mM Tris, 100 mM NaCl, pH 8.5 was concentrated to 5 mg/ml. For both, CryoEM and crystallography, the enzyme fractions were activated using 10 mM NaHCO_3_ in the presence of a 2-fold molar excess of CABP for about 30 minutes.

### Radiometric carboxylation assays

Time curve assays: The assays were adapted from Schulz et al.^66^ Prior to each assay, protein stock concentration was measured via BCA assay (Thermo Scientific, MA) and the condensate formation was assessed via light microscopy (supplementary figure 2) under reaction conditions. Reactions were conducted with NaH^14^CO_3_ or NH_4_H^14^CO_3_ stocks with specific radioactive activities between 4 to 8 mCi/mmol. The solubility of CO_2_ in water (mol/L/atm) was calculated using Henderson-Hasselbalch equation using previously published values^67^. Time curve assays were performed aerobically in 50-100 μL volumes in 1.5 mL crimp neck amber vials (VWR) at 25°C and 300 rpm using a thermomixer pro (Cellmedia, Germany). For each data set, 4 time points with each 3 replicates for both CbbM and CbbM-IDP in the same shaker were measured. To this end, 1 μM CbbM/CbbM-IDP, 0.1 mg/ml CA, 1 mM MgCl_2_ was activated for 5 minutes using 5 mM ^14^CO_3_^2-^ stocks in 50 mM Tris, pH 7.5-8.0 (depending on condensate stability). Subsequently, 0.3 mM RuBP was added to start the reaction. After 2, 10, 30 and 90 min the reactions were quenched using 10% final v/v of 50 % v/v formic acid. Reactions were left to evaporate on a heating block at 95 °C ON. Residual material was re-suspended in 500 μL ddH_2_O, transferred to 5 mL scintillation vials, and mixed with 4.5 mL scintillation cocktail (ROTISZINT®HighCapacity, Carl Roth). The amount of acid-stable radioactivity was quantified using a Beckman LS 6000 scintillation counter. Initial rates were computed using the 2 min time point and normalized with the respective active site content. Active site content was quantified via [^14^C]-CABP binding. To this end, the same reaction mixtures with cold CO_3_^2-^ and CABP instead of RuBP were incubated in the reaction buffer supplemented with 500 mM NaCl to prevent condensate formation for 1 hour. The mixtures were centrifuged for 10 min at 10000xg. Subsequently, 35 μL supernatant was applied to a SERAgel Pro 250 column (ISERA, Germany) on a Agilent 1200 HPLC system and isocratically eluted in the same buffer. Fractions containing the protein were collected (2 mL total volume), mixed with 12 mL scintillation cocktail and measured on the scintillation counter. Active site content was computed from the known counts/concentration of the [^14^C]-CABP stocks. For total carboxylation, the ratios of the maximally reached scintillation count in the time curves of Rubisco and Rubisco-fusion were used. If necessary, multiple data sets were combined by first computing the ratios of individual data sets and then averaging the ratios to account for fluctuations in assays. Errors were propagated to the ratios and, subsequently, to the final ratio via empirical error propagation. In case of anaerobic initial rate comparisons, assay buffers, vials and stocks were degassed with CO_2_ free nitrogen for at least 3 hours.

^3^H specificity assays: The assays were performed exactly as reported in Schulz et al.^66^ Briefly, CbbM and CbbM-IDPs were incubated in 20-mL septum capped glass scintillation vials containing 1 mL 50 mM Tris, pH 7.5-8.0, 1 mM MgCl_2_, and 0.1 mg/mL carbonic anhydrase. Reaction mixtures were equilibrated in defined gas mixtures containing 995009 ppm O_2_ and 4991 ppm CO_2_ (Air Liquide, Germany) before initiating the reactions by the addition of 1 mM [1-3H]-RuBP. After 1 hour, products were dephosphorylated using alkaline phosphatase (10 U/reaction) for another 30 min, and, subsequently separated on an HPX-87H column (Bio-Rad, CA). ^3^H-glycerate and ^3^H-glycolate peaks were quantified using liquid scintillation counting. The specificity factor S_C/O_ was calculated as described previously^47^. Reaction buffers were adapted to condensate forming conditions using the same buffers as for the time curve assays.

### LC-MS/MS specificity assays

Assays were adapted from Schulz et al.^66^: Buffers and vials were degassed with CO_2_ free nitrogen and equilibrated in 995009 ppm O_2_ and 4991 ppm CO_2_ (Air Liquide, Germany) prior to reaction start and during reaction. Reactions were run in 0.5 mL volumes in 1.5 mL crimp neck amber vials (VWR) equipped with gas tight conical rubber stoppers in a thermomixer pro (CellMedia, Germany) at 25 °C. For each reaction, 1 μM CbbM and CbbM-IDPs were used. Reactions were supplemented with 0.1 mg/mL CA and then activated for 5 min before addition of 1 mM RuBP. Reactions were run in quadruplicates for 30 min and quenched using 50 μL formic acid. Subsequently, 35 μL were centrifuged for 2 min at 3000xg and submitted for LC-MS/MS measurement.

### LC-MS/MS method

Quantitative determination of RuBP, 2-PG and 3-PG was performed using a LC-MS/MS. The chromatographic separation was performed on an Agilent Infinity II 1290 HPLC system using a SeQuant ZIC-pHILIC column (150 × 2.1 mm, 5 μm particle size, peek coated, Merck) connected to a guard column of similar specificity (20 × 2.1 mm, 5 μm particle size, Phenomoenex) a constant flow rate of 0.1 ml/min with mobile phase A with mobile phase comprised of 10 mM ammonium acetate in water, pH 9, supplemented with medronic acid to a final concentration of 5 μM (A) and 10 mM ammonium acetate in 90:10 acetonitrile to water, pH 9, supplemented with medronic acid to a final concentration of 5 μM (B) at 40° C . The injection volume was 0.5 μl. The mobile phase profile consisted of the following steps and linear gradients: 0 – 1 min constant at 75 % B; 1 – 6 min from 75 to 40 % B; 6 to 9 min constant at 40 % B; 9 – 9.1 min from 40 to 75 % B; 9.1 to 20 min constant at 75 % B. An Agilent 6470 mass spectrometer was used in negative mode with an electrospray ionization source and the following conditions: ESI spray voltage 2000 V, nozzle voltage 1000 V, sheath gas 300° C at 12 l/min, nebulizer pressure 20 psig and drying gas 150° C at 11 l/min. Compounds were identified based on their mass transition and retention time compared to standards. Chromatograms were integrated using MassHunter software (Agilent, Santa Clara, CA, USA). Absolute concentrations were calculated via standard addition, spiking each sample with a mixed stock solution of the target analytes at different concentrations (0 μM, 10 μM, 25 μM, 50 μM, 100 μM and 250 μM) and analysing each sample six times. Mass transitions, collision energies, Cell accelerator voltages, and Dwell times have been optimized using chemically pure standards. The parameter settings of all targets are given in table 1.

**Table 1.** Parameter settings for the Agilent 6470 mass spectrometer.

| Name | Precursor Ion | Product Ion | Collision energy [V] | Fragmentor Voltage [V] | Cell Accelerator Voltage [V] | Dwell time [msec] | Polarity |
| --- | --- | --- | --- | --- | --- | --- | --- |
| RuBP | 309.09 | 97 | 17 | 140 | 5 | 90 | Negative |
| RuBP | 309.09 | 79 | 59 | 140 | 5 | 90 | Negative |
| Ru5P | 229 | 97 | 8 | 140 | 5 | 90 | Negative |
| Ru5P | 229 | 79 | 48 | 140 | 5 | 90 | Negative |
| 3-PG | 185.06 | 85.06 | 0 | 140 | 5 | 90 | Negative |
| 3-PG | 185.06 | 97 | 13 | 140 | 5 | 90 | Negative |
| 3-PG | 185.06 | 79 | 32 | 140 | 5 | 90 | Negative |
| 2-PG | 155 | 97 | 19 | 140 | 5 | 90 | Negative |
| 2-PG | 155 | 79 | 33 | 140 | 5 | 90 | Negative |

### Crystallisation and x-ray structure determination

Crystallization was performed by sitting-drop vapor-diffusion at 16 °C. Purified inactive CbbM in 20 mM Tris buffer (pH 8.5), containing 100 mM NaCl was mixed with condition comprised of 1.2 M ammonium chloride, 100 mM MES (pH 6), 20 %(w/v) PEG6000 in a 1:1 ratio with a final drop volume of 1 µL. The mother liquor was supplemented with 30% (v/v) PEG200, before the crystals were plunge frozen in liquid nitrogen. Crystallization of activated CbbM in buffer containing 20 mM Tris (pH 8.5), 100 mM NaCl, 0.2 mM CABP, and 10 mM sodium bicarbonate was mixed with a condition containing 200 mM Bis-Tris propane (pH 7.3) and 20 % (w/v) PEG4000. Prior to plung freezing the crystals in liquid nitrogen the mother liquor was supplemented with 33 % (v/v) PEG200.

X-ray diffraction data (supplementary table 5) were collected at the PETRA III beamline P13 of DESY (Deutsches Elektronen-Synchrotron, Hamburg). Data were processed with XDS^68^. Structures were solved by molecular replacement using Phaser of the Phenix software package (v1.21)^69^ using an AlphaFold2^70^ model as search template and refined with Phenix.Refine. Manual modelling, refinement, and ligand fitting, as well as water picking was performed in Coot (v.0.9.8.3)^71^. Final positional and B-factor refinements were performed using Phenix.Refine. Structural models for the inactive form and activated CABP bound form were deposited to the Protein Data Bank in Europe (PDBe) under PDB accession 8S21 and 8S20, respectively (supplementary table 5). Figures were made using PyMOL 2.5 (The PyMOL Molecular Graphics System, Version 2.5 Schrödinger, LLC. (www.pymol.org)).

### cryoEM sample preparation, data acquisition and single particle reconstruction

The CbbM-EPYC1Δα stock was activated using 10 mM NH_4_HCO_3_ and a 2-fold excess of CABP for 30 min prior to sample preparation. Subsequently, 2.7 μL 50 mM Tris, 0 mM NaCl, pH 7.5 was pipetted onto a C-Flat R2/1 300 copper mesh grids that were glow-discharged for 45 s immediately before use and 0.3 μL of the protein stock was added to the drop resulting in ∼1 μM final protein concentration.

The sample was blotted for 3 s with blot force 20 at 100% relative humidity and 4 °C using a Vitrobot Mark IV (Thermo Scientific). Grids were plunge frozen in liquid ethane cooled by liquid nitrogen, clipped into Autogrid cartridges, and used for data collection immediately.

CryoEM data were acquired on a Titan Krios G3i electron microscope (Thermo Scientific), operated at an acceleration voltage of 300 kV and equipped with a BioQuantum-K3 imaging filter (Gatan). Data were measured in electron counting mode at a nominal magnification of 130,000× (0.655 Å/pixel) with a total dose of 60 e^–^/A^2^ (60 fractions), using the aberration-free image-shift (AFIS) correction in EPU (Thermo Scientific). Five images were acquired per foil hole, and the nominal defocus range for data collection was - 1.2 to - 2.4 μm. Foil holes were selected based on capturing a sufficiently thin portion of the condensates around the too dense parts.

Micrographs were processed in cryoSPARC (Structura Biotechnology Inc., Canada, v4.4.1). First 11525 micrographs got motion and CTF corrected, then low quality micrographs were excluded yielding 7116 remaining micrographs. 45233 particles were manually picked since the small size of the structured domain precluded reliable automated picking in cryoSPARC. Through several classification steps the final structure reconstruction was done with 18706 high quality particles. Detailed SPR is shown in supplementary figure 14. A CryoEM map fitting was initially performed in UCSF-ChimeraX (v1.7)^72^ using an AlphaFold2^70^ model as template. The resulting model was manually refined in Coot (v0.9.8.3)^71^. Automatic refinement of the structure was performed using phenix.real_space_refine of the Phenix (v1.21)^69^ software suite. The resulting model and electron density map were deposited in the Protein Data Bank in Europe and the Electron Microscopy Data Bank with the accession codes 8S22 and EMD-19652, respectively. Model statistics are listed in supplementary table 6.

### Confocal microscopy for fluorescence detection of carboxylation, polarity estimation and carbonic anhydrase partitioning

The activity in the condensates was qualitatively observed using a fluorescence coupled assay (supplementary figure 6). The precise concentrations are displayed in table 2. Pgmi was produced in house, HRP acquired from Sigma (P8375), enolase from Sigma (E6126), pyruvate kinase from Roche (10109045001), carbonic anhydrase from Sigma (C2624), and pyruvate oxidase from Sigma (P4591). First, all components except the Rubisco and the RuBP are pipetted into 384 glass bottom plates (Cellvis, 384 well glass bottom plate P384-1.5H-N). Then the protein is added and the mixture is checked for condensate formation. After Rubisco activation for several minutes the RuBP is added to observe the increase in fluorescence in a confocal epi-fluorescence microscope (Leica TCS SP8 X with a Leica DFC500 camera). Images were recorded at 1024x1024 pixels with 80.17 nm pixel size and a rate of 0.644 images/s for 25 minutes. Samples were excited at 2 % LP at 570 nm and recorded with a 585-600 nm detection window.

**Table 2:** Reaction mixtures for the fluorescence detection of Rubisco carboxylation activity.

| 40 x Mix | Volumes | Stock concentrations | Final concentrations |
| --- | --- | --- | --- |
| Kpi buffer | 82.26415 μL | pH 7.5, 50 mM | |
| Amplex | 25 μL | 20 mM | 2.5 mM |
| Pgmi | 37.73585 μL | 265 μM | 50 μM |
| HRP | 2 μL | 1000 U/mL | 10 U/mL |
| ADP | 20 μL | 500 mM | 50 mM |
| Enolase | 10 μL | 5 mg/mL | 0.25 mg/mL |
| Carbonic anhydrase | 10 μL | 2 mg/mL | 0.1 mg/mL |
| Pyruvate kinase | 10 μL | 25 U/mL | 1.25 U/mL |
| V(total) | 200 μL | | |

| Reaction mix | Volumes | Stock concentrations | Final concentrations |
| --- | --- | --- | --- |
| Rubisco | x | | 1 μM |
| 40 x Mix | 1 μL | | |
| MgCl<sub>2</sub> | 0.5 μL | 0.2 M | 2.5 mM |
| carbonate | 0.5 μL | 500 mM | 6.25 mM |
| Pyruvate oxidase | 1 μL | 1 U/mL | 0.025 U/mL |
| 1-5-Ribulose biphosphate | 0.4 μL | 100 mM | 1 mM |
| Tris buffer | Rest | pH 7.5, 50 mM |  |
| V(total) | 40 μL | | |

Polarity of condensates was estimated using the assay reported in Küffner et al^42^. Condensates were formed with 1 μM of fusion in 50 mM Tris, 0 mM NaCl and pH 7.5. Images were recorded at 1024x1024 pixels with 41.01 nm pixel size and a rate of 0.375 images/s. Samples were excited at 2 % LP with 405 nm and detected in a window from 425 nm to 784 nm with a 5 nm bandwith and 5 nm steps. Subsequently the intensities were extracted and a gamma distribution fitted to the data to estimate the maximum intensity wavelength. Polarity was inferred by a standard curve using reported literature data (supplementary figure 10)^73,74^.

Carbonic anhydrase partitioning was assessed using Atto425 tagged carbonic anhydrase. To his end, we incubated 2 mg/mL carbonic anhydrase from bovine erythrocytes (Sigma-Aldrich) with a ∼2-fold molar excess of Atto425-Malemide (ATTO-TEC) for 12 h on ice. Subsequently, we separated the tagged protein from unreacted Atto425 with a 10/30 Superdex 200 pg increase (Cytiva) isocratically in 50 mM Tris, 500 mM NaCl, 2 mM 2-BME, pH 8.5. The tagged protein was concentrated using 500 μL Amicon 10 kDa spin filters. Condensates were formed using 1 μM of fusion protein in 50 mM Tris, 0 mM NaCl, pH 8.0. After 5-10 min a mixture of 10% CA-ATTO425 and 90% CA with a total final concentration of 0.1 mg/mL was added to the condensates. After a couple of minutes, images were recorded at 1024x1024 pixels with 80.17 nm pixel size, a rate of 0.241 images/s and a frame averaging of 8. Samples were excited at 2 % LP with 405 nm and recorded with a 465-505 nm detection window.

### Mass photometry size analysis

Microscope coverslips (1.5 H, 24 x 60 mm, Carl Roth) and CultureWell^TM^ Reusable Gaskets (CW-50R-1.0, 50-3mm diameter x 1 mm depth) were three times cleaned with MQ-H_2_O and isopropanol followed by drying under a stream of pressurized air. Gaskets were dried further at RT overnight.

Immersion oil (Immersion Oil 518 F, Zeiss, Germany) was applied to the MP objective and, subsequently, dry gaskets were assembled on coverslips and placed on the stage of an One^MP^ mass photometer (MP, Refeyn Ltd, Oxford, UK). Then 18 μL of SEC-buffer was applied to the gasket wells. After focusing, 2 μL sample was added, rapidly and measurements were started (<15 s after sample addition). Data was acquired for 60 s at 100 frames per second using Acquire^MP^ (Refeyn Ltd, v1.2.1). Samples were prepared by diluting purified protein to 0.25 μM concentration in 50 mM Tris, 500 mM NaCl, 10 mM MgCl_2_, pH 8.5. MP contrast values were calibrated to molecular masses using commercial NativeMark^TM^ Unstained Protein Standard (Thermo Fisher, MA). The instrument was calibrated using a 30-fold dilution of NativeMark, which was diluted further 10x on the instrument. Acquired images were processed using DiscoverMP (Refeyn Ltd, v.1.2.3).

### Mass action and mass transfer model simulations and data fit

Differential equation systems were implemented in python 3.10 and solved using scikits.odes based on the SUNDIALS suit^53–55^. Data was fitted using maximum likelihood estimators and globally optimised using differential evolution algorithm from scipy.optimize^75,76^. The differential equation system for Rubisco was adapted from previous simulation work shown below^41^:

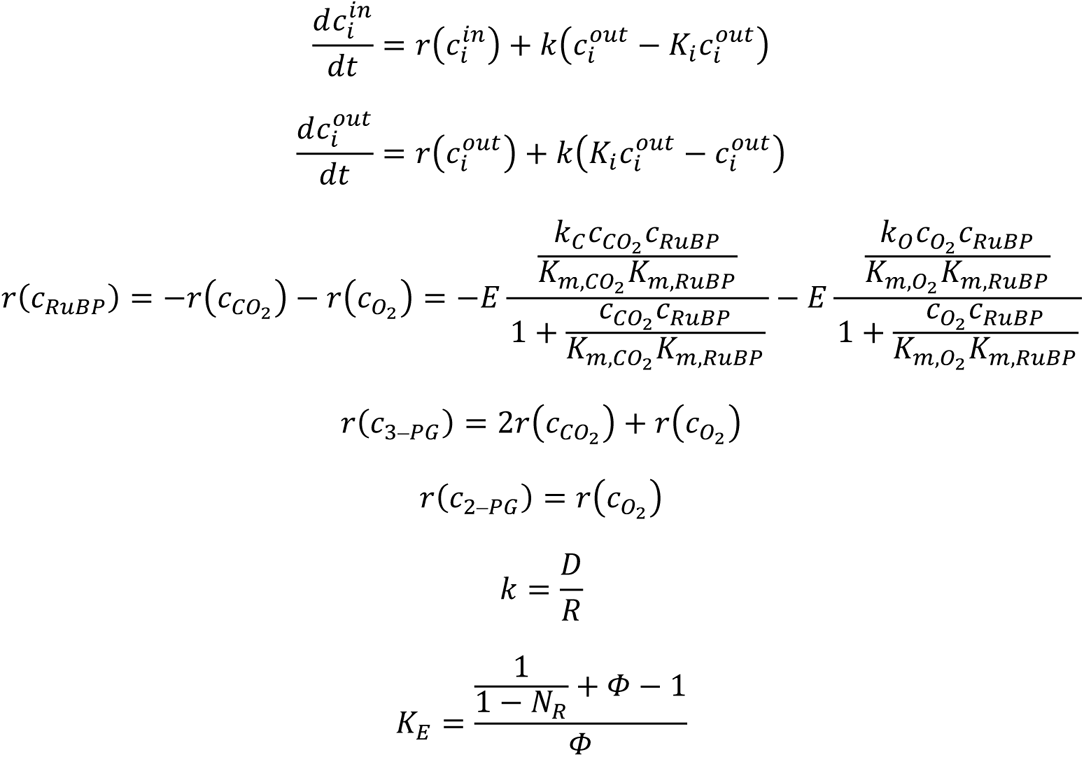

Where 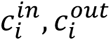 denote the concentrations of compound 𝑖 inside and outside of the condensates, 𝑘, 𝐾_𝑖_ denote the mass transfer coefficient and the partition coefficient of the respective compound, 𝑟(𝑐_𝑖_) denotes the rate of the respective compound, 𝐸 denotes the active site concentration, 𝑘_𝐶_ , 𝑘_𝑂_denote the catalytic constant of carboxylation and oxygenation respectively, and 𝐾_𝑚,𝐶𝑂_2__ , 𝐾_𝑚,𝑂_2__ , 𝐾_𝑚,𝑅𝑢𝐵𝑃_ denote the michaelis constant of CO_2_, O_2_, and RuBP respectively. The diffusion coefficient (D) was assumed to be 10 μm^2^/s based on small molecule diffusion across condensates^41^, the average condensate radius (R) was assumed to be 10 μm at a phase volume fraction Φ of 1% based on estimations from previous measurements^41^. The average radius was assumed to linearly scale with the phase volume fraction in a first approximation.

