## Supplementary Information for "Bottom-up reconstruction of minimal pyrenoids provides insights into the evolution and mechanisms of carbon concentration by EPYC1 proteins"

Supplementary Figures 1-13

Supplementary Tables 1-6

**Supplementary figure 1 SDS-PAGE gels of investigated proteins.** A.) SDS-PAGE gel of CbbM and CbbM-IDP fusion proteins from the EPYC1 homologs and deletion mutants. All homologs are ΔCtp (chloroplast targeting protein) to assure correct localisation to the cytosol for recombinant purification. B.) SDS-PAGE gel of CbbM and CbbM-IDP fusion of EPYC1 ancestors.

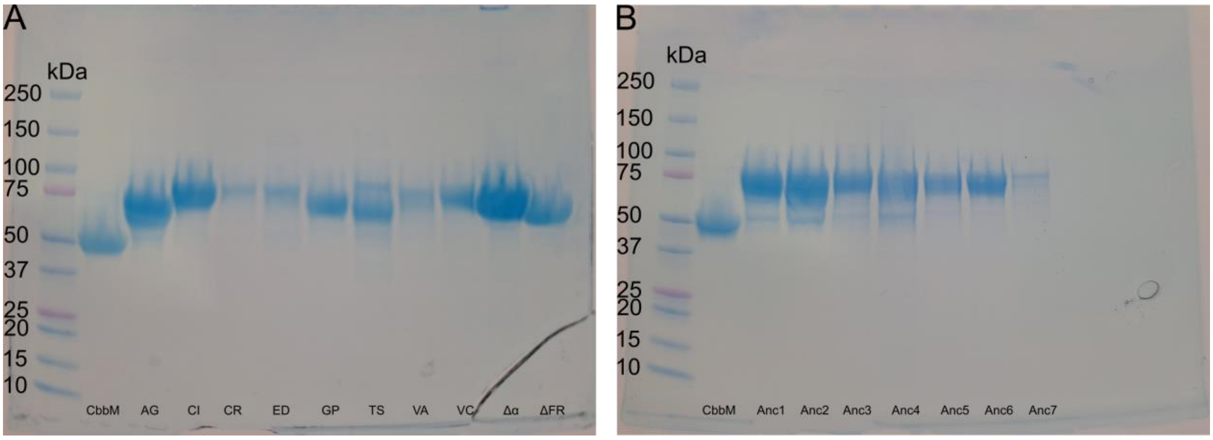

**Supplementary figure 2 Microscopy images of CbbM-IDP fusions under reaction conditions.** Microscopy images of CbbM-IDP fusion proteins at 1 µM concentration, 0.1 mg/mL CA, 50 mM Tris, pH 7.5-8.0, 5 mM $NaHCO_3$ (Δα, CR, CI) or $NH_4CO_3$ (all ancestors, ED, GP ,TS, VA, VC).

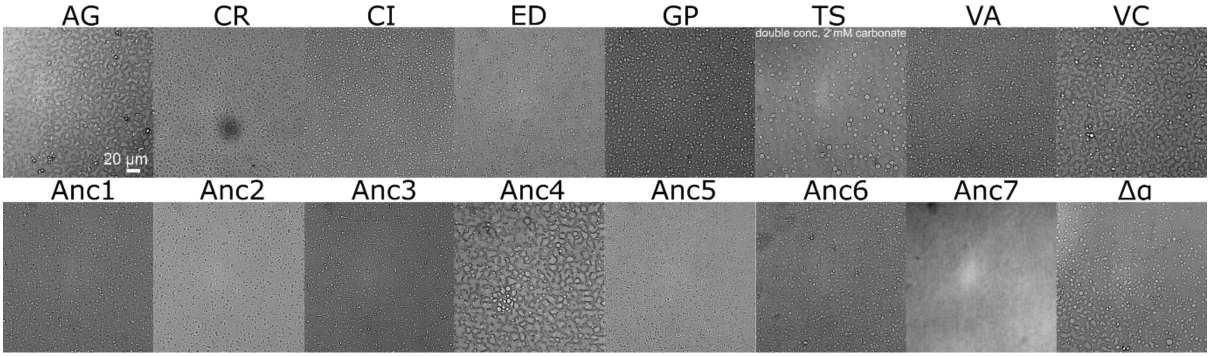

**Supplementary figure 3 Microscopy images of CbbM-EPYC1$_{CR}$Δα and CbbM-Laf1.** Transmission microscopy images of condensates (CbbM-EPYC1$_{CR}$Δα) and aggregates (CbbM-Laf1N) at 1 µM protein concentration in 50 mM Tris, 0 mM NaCl, pH 8.0. CbbM- EPYC1$_{CR}$Δα forms condensates, while CbbM-Laf1N shows solidified aggregates.

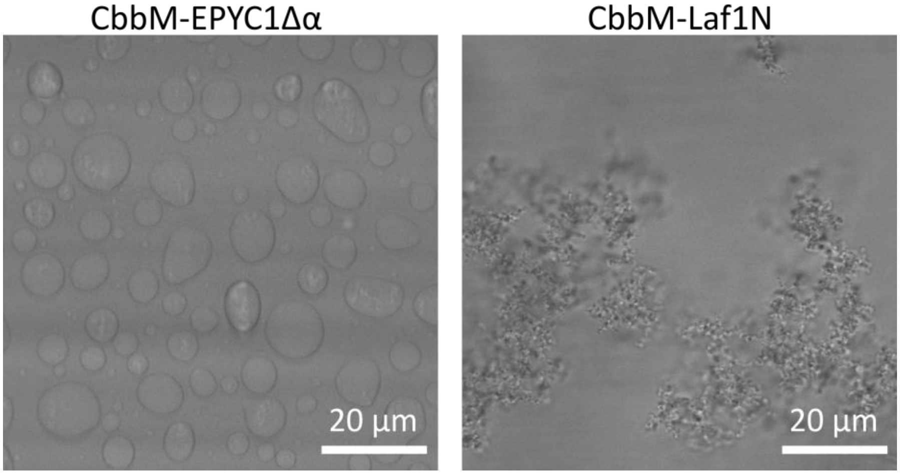

**Supplementary figure 4 Logo plots indicating amino acid composition in different non-EPYC related IDPs.** Sequences were extracted from literature indicating clear LLPS behaviour of the respective IDPs[4,10,26–33].

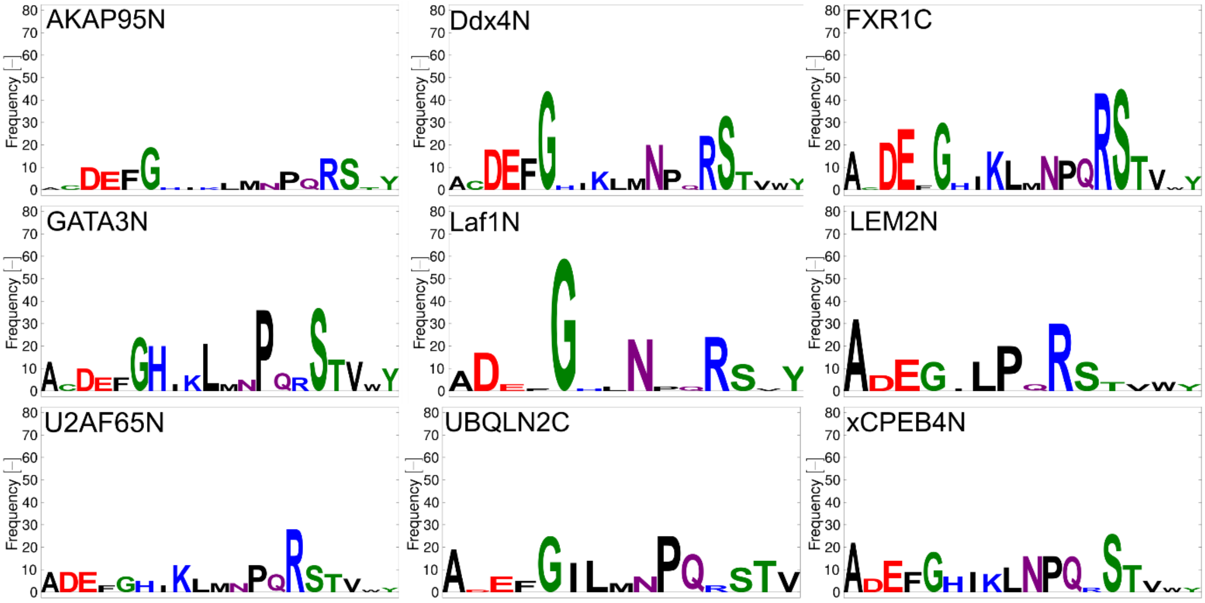

**Supplementary figure 5 Logo plots indicating amino acid composition in different EPYC homologues of the EPYC1 from *C. reinhardtii*.** Sequences were blasted using NCBI protein BLAST at the 06.09.2022.

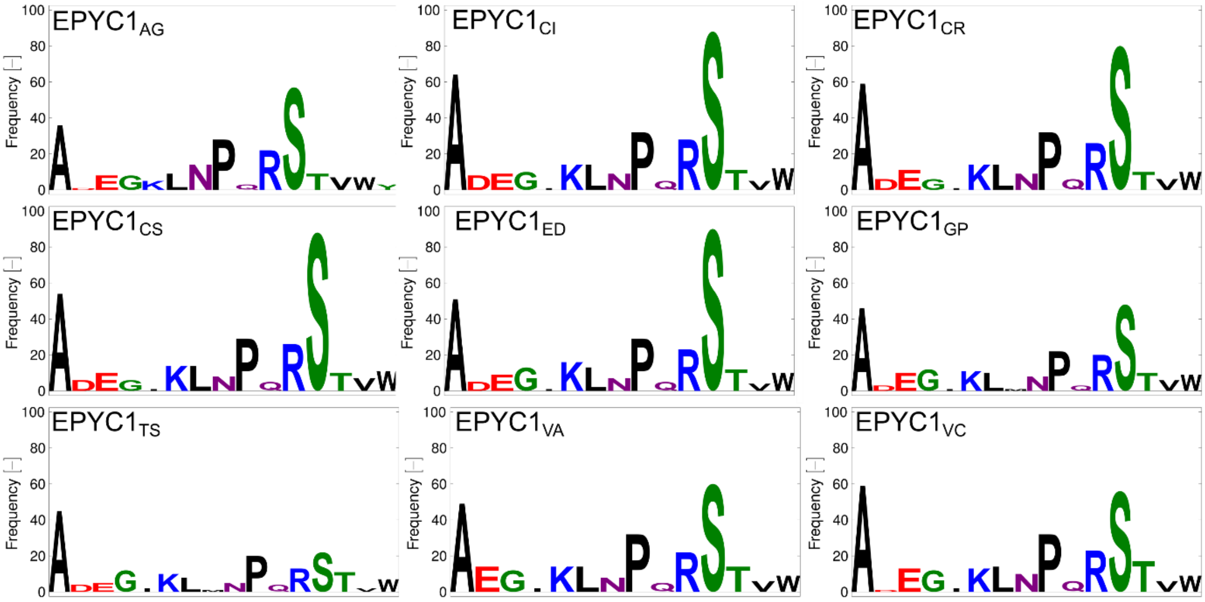

**Supplementary figure 6 Scheme describing the fluorescence Rubisco carboxylation activity assay.** RuBP can be carboxylated yielding 3-PG or oxygenated yielding 2-PGA by Rubisco. Phosphoglycerate mutase shifts the phosphate group from position 3 to 2 on 3-PG. Using an enolase the glycerate can be rearranged yielding phosphoenol pyruvate, which subsequently gets dephosphorylated via a pyruvate kinase. Pyruvate is finally oxidized to acetyl phosphate and hydrogen peroxide catalysed by pyruvate oxidase. Hydrogen peroxide evolution is detected using horseradish peroxidase and Amplex Red.

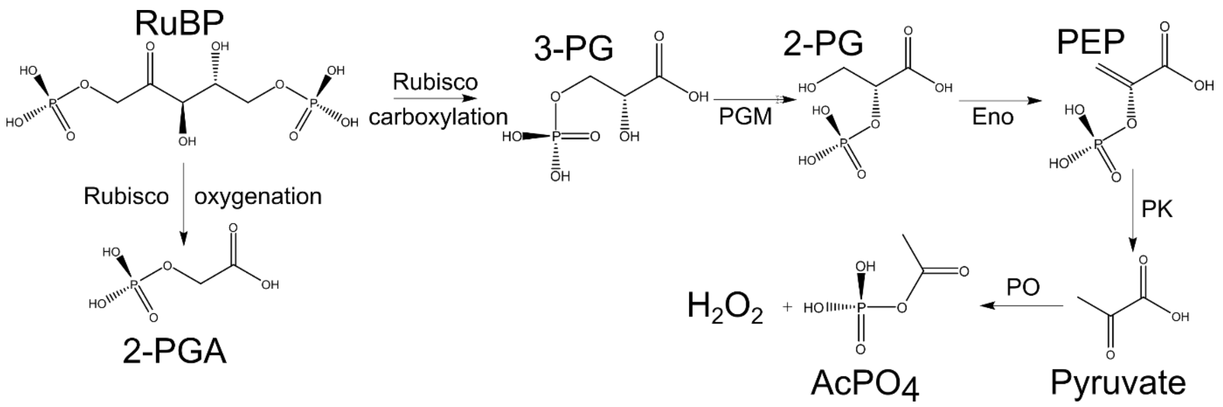

**Supplementary figure 7 Protein and species tree of the EPYC homologues and respective species.** A.) The IQ-TREE result from ASR relating the EPYC homologues to each other is shown[58]. B.)The species tree of the EPYC homologues containing *Chlamydomonadales* species from timetree.org is shown[34]. The species tree was used to root the protein tree to the oldest evolutionary node. Additionally, age, oxygen and carbon dioxide data in relation to the species tree is depicted.

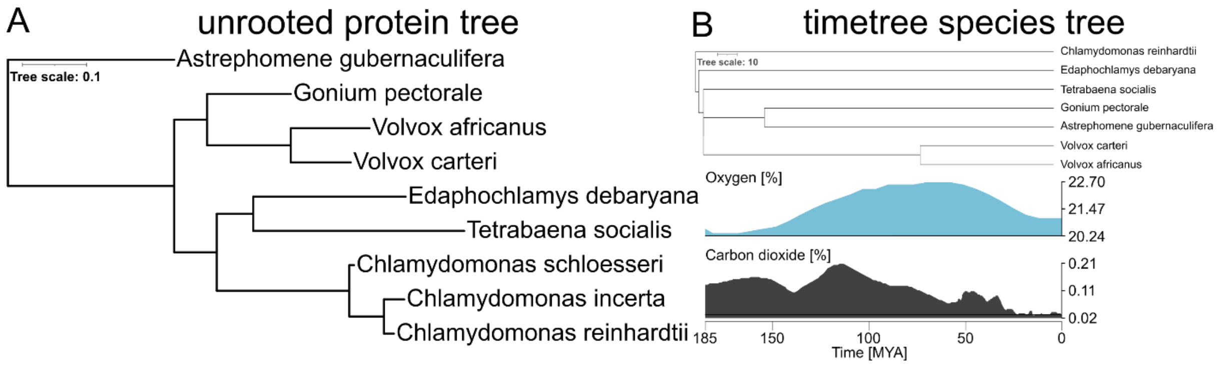

**Supplementary figure 8** **CbbM-EPYC1$_{CR}$ condensates under different reaction conditions** A.) Brightfield microscopy images of the condensates formed by CbbM-EPYC1$_{CR}$ using 1 µM protein in 50 mM Tris, pH 7.5, 0.1 mg/mL CA, 5 mM $NaHCO_3$, 0.3 mM RuBP, 10 mM $NaHCO_3$ (B), 0.3 mM RuBP (C; different experiment to A) and 1 mM RuBP.

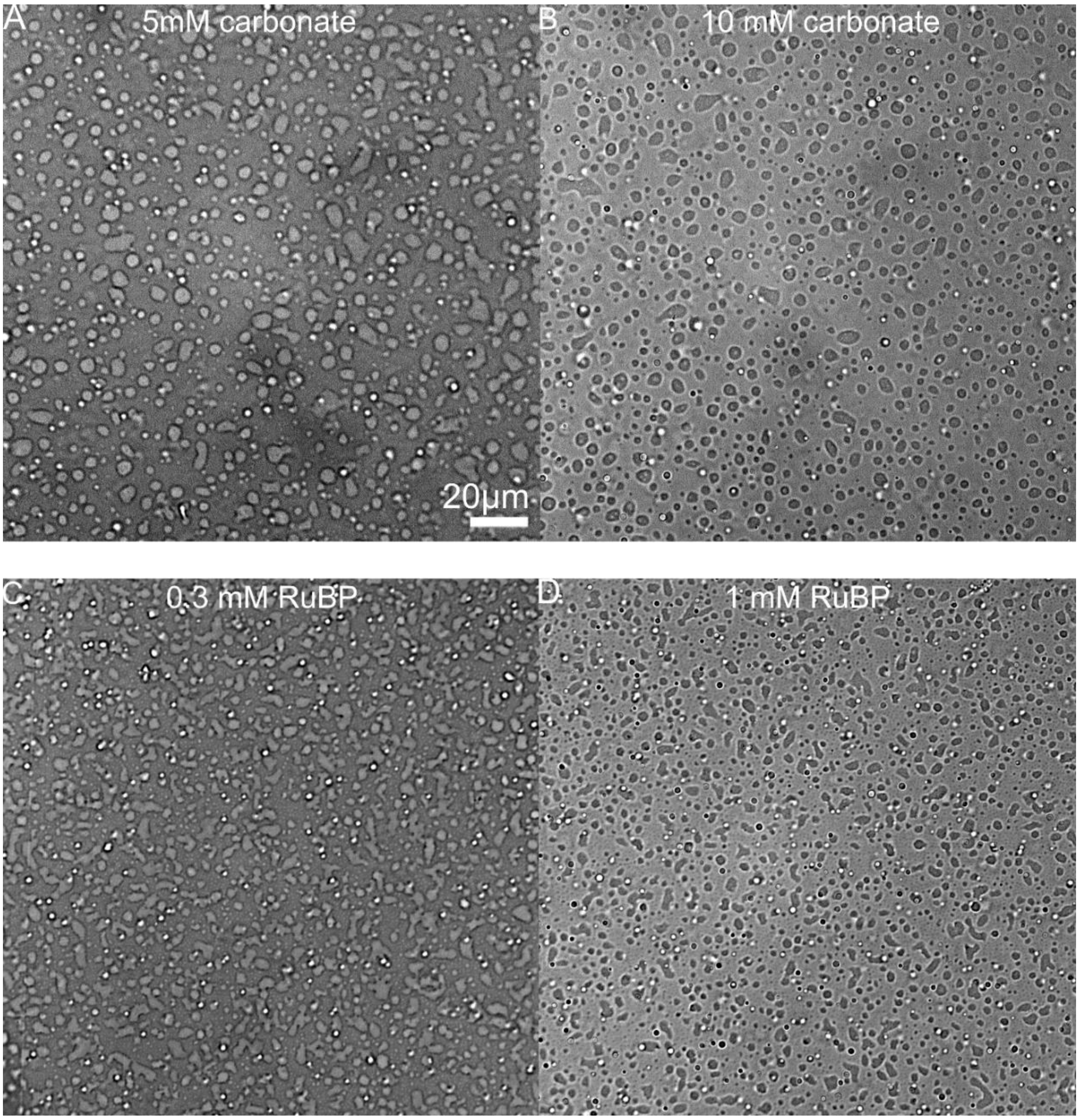

**Supplementary figure 8 PRODAN polarity estimation of condensates formed by EPYC1 variants.** A.) Exemplary fluorescence confocal microscope images of the condensates formed by $EPYC1_{CR}\Delta\alpha$ and by CbbM-$EPYC1_{CR}\Delta\alpha$ indicating the different maximum fluorescence emission colour. Condensates were formed in 50 mM Tris, pH 8.0 using 1 µM protein (10 µM for $EPYC1_{CR}\Delta\alpha$) and ~1mM of PRODAN. B.) Plot showing the standard curve using literature data[73,74] on emission maxima in different solvents and the inferred apparent polarities of the condensates formed by selected EPYC1 homologues.

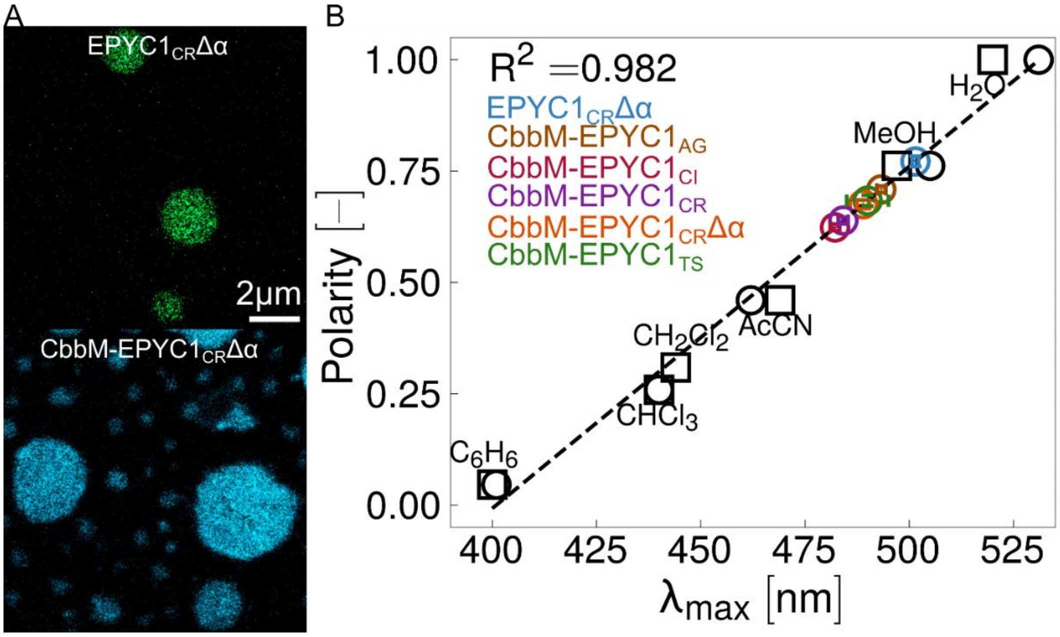

**Supplementary figure 9 Carbonic anhydrase partitioning into condensates.** Condensates were formed in pH 7.5-pH 8.0 50 mM Tris, 0 mM NaCl, ~0.1 mg/ml CA with 10% (stock v/v) ATTO-425 tagged carbonic anhydrase. At least two biological replicates, with each at least four images were recorded. A.) Exemplary images of different EPYC1s and CbbM-EPYC1 fusions are shown. B.) Fluorescence intensity ratios as a proxy for CA partitioning ($K_{C/S}$) were calculated by measuring inside and outside the condensates in an equally distributed manner. Ratio errors were calculated form the standard deviation of fluorescence intensity inside and outside the condensates and propagated to the ratio. P-values were calculated against a ratio of 1 indicating no partitioning in either direction using a two sided t-test (p<0.05=*, <0.01=**, <0.001=***, <0.0001=****).

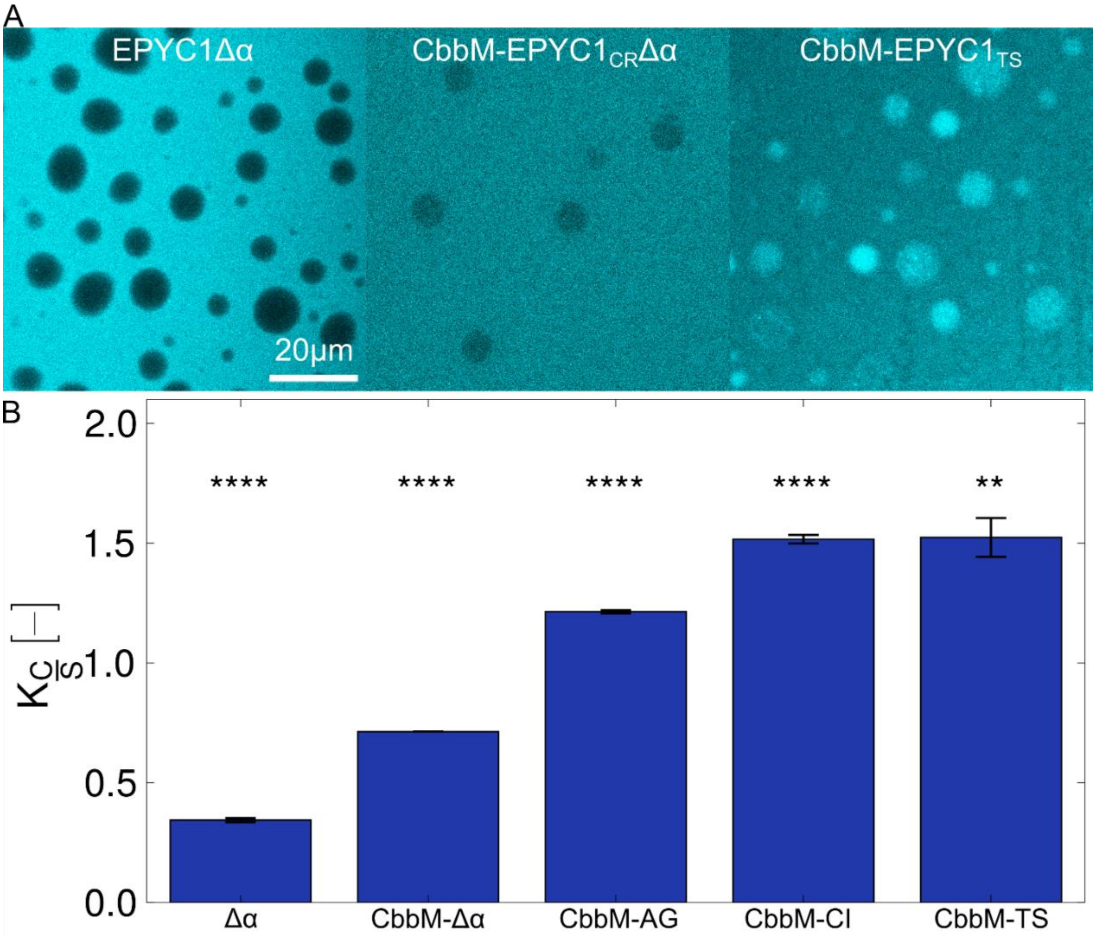

**Supplementary figure 10 Fibrillar structures emerging in 2D classification of CbbM-EPYC1$_{CR}$Δα at the edges of the condensates.** A.) Exemplary 2D-classes emerging after 2D classification showing fibrillary elongated structures pointing towards EPYC1$_{CR}$Δα IDPs or bundles of them. b.) *Ab initio* volume reconstruction from the 2D-classes showing elongated volume in the size range of several IDPs bundled up.

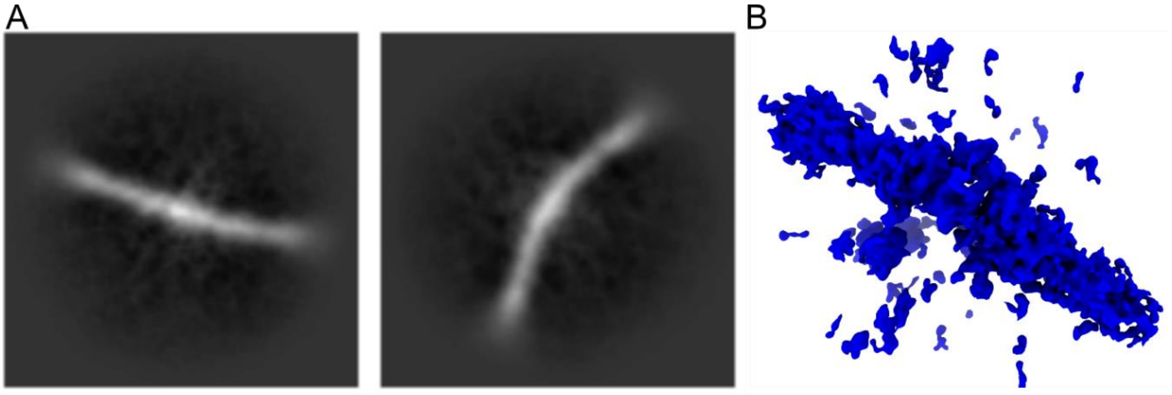

**Supplementary figure 11 Comparison of CbbM-EPYC1$_{CR}$Δα and CbbM-EPYC1$_{CR}$.** A.) Brightfield microscopy images of the condensates formed by the α-helical deletion mutant of EPYC1 (Left) and the WT EPYC1$_{CR}$ (right) using 1 µM protein in 50 mM Tris, pH 8.0, 0.1 mg/mL CA, 5 mM $NaHCO_3$, 0.3 mM RuBP. B.) Corresponding radiometric measurements of carboxylation under aerobic conditions using 0.3 mM RuBP, 5 mM $CO_3^{2-}$, 0.1 mg/mL CA at 25°C.

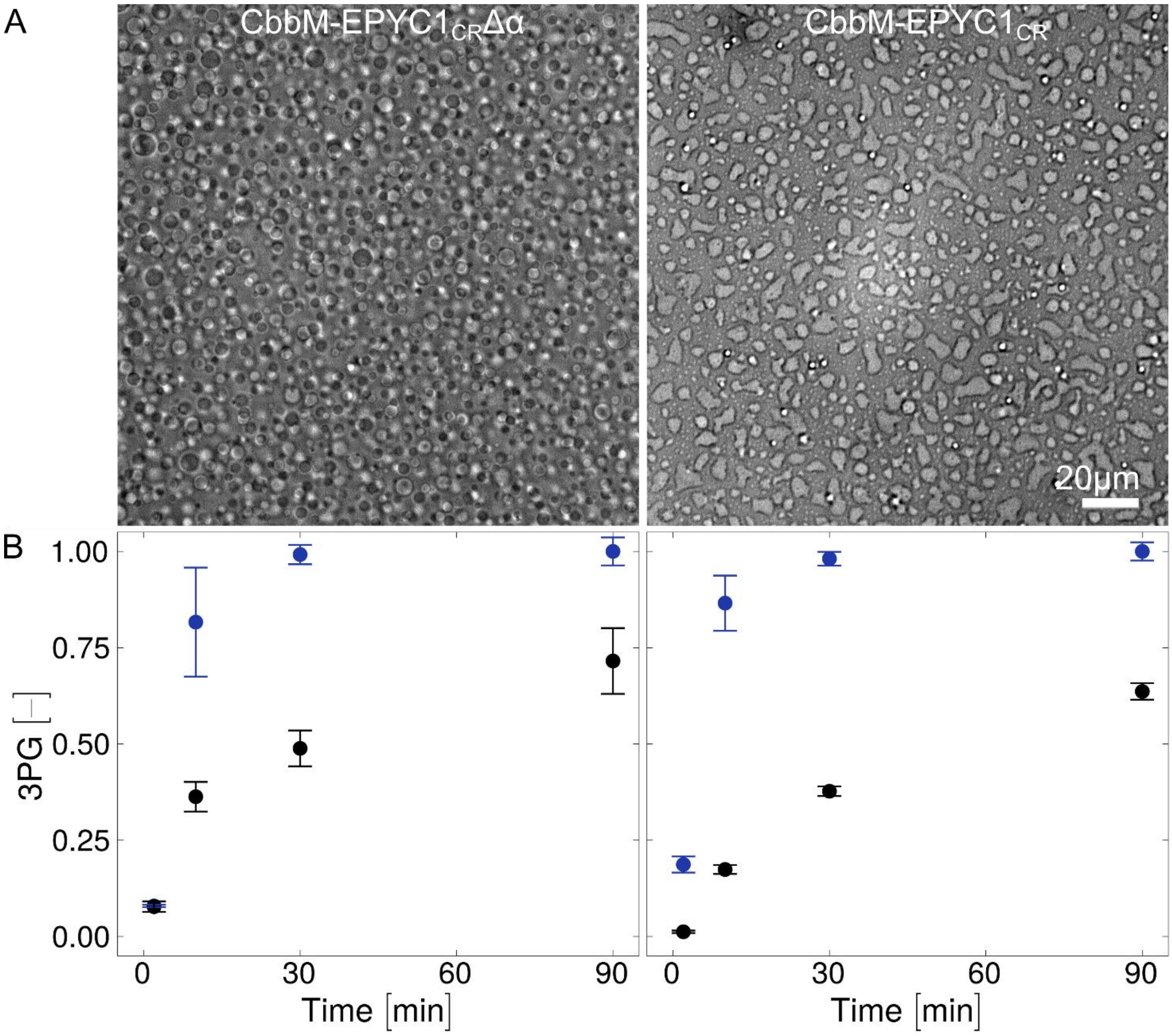

**Supplementary figure 12 Mass photometry data of solvated CbbM and CbbM-EPYC1$_{CR}$Δα.** A.) Mass photometry of 25 nM CbbM in 50 mM Tris, pH 8.0, 500 mM NaCl. CbbM is visible as dimer (~100 kDa) and as monomer (~50 kDa). B.) Mass photometry of 25 nM CbbM -EPYC1Δα in 50 mM Tris, pH 8.0, 500 mM NaCl. CbbM -EPYC1Δα is visible as dimer (~150 kDa) and monomer (~75 kDa). The CbbM -EPYC1Δα has very low detachment events (negative range), indicating the stickiness of CbbM -EPYC1$_{CR}$Δα on the surface of the cover-slide in contrast to CbbM.

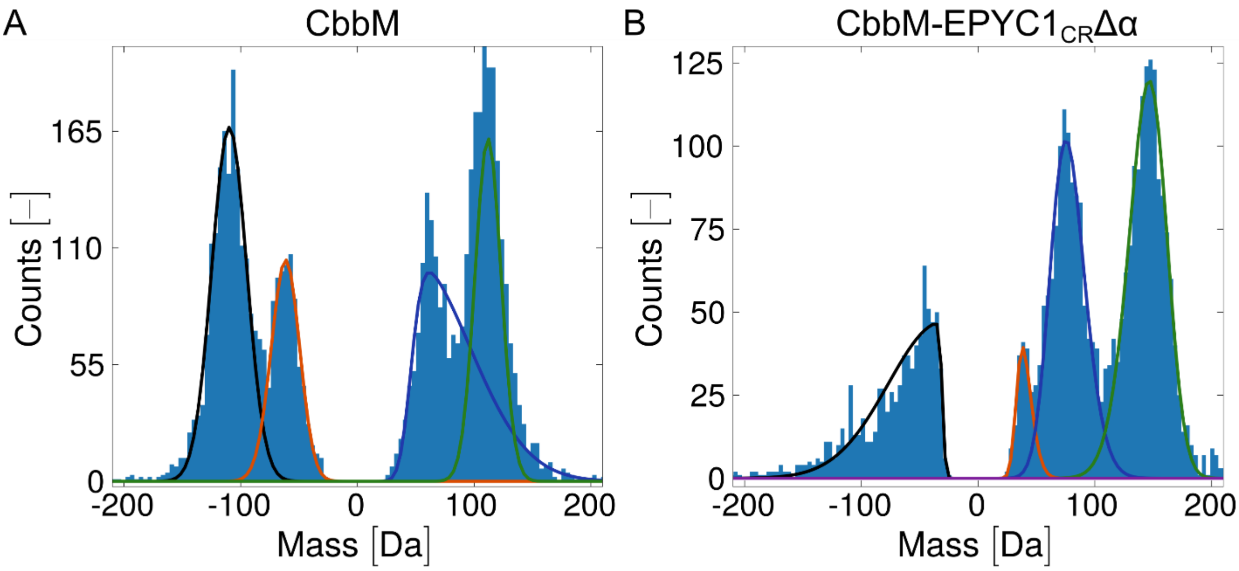

**Supplementary figure 13 Cryogenic electron transmission microscopy single particle reconstruction workflow inside the condensates.** A.) Flowchart depicting the CryoEM SPR workflow in cryoSPARC. B.) Density map of the largest class (18706 particles) from the *ab initio* reconstruction from 29247 particles is depicted. V.) Final density map of CbbM from the CbbM-EPYC1$_{CR}$Δα inside the condensates after non-uniform refinement and local refinement is shown. D.) Local resolution estimate map of CbbM from C.) is illustrated. E.) The GSFCS plot from the final local refinement is presented indicating a final global resolution of 3.41 Å. F.) Guinier plot of the local refinement is shown indicating a B-factor of 98.7. G.) Viewing direction distribution plot of the local refinement is depicted indicating an overall uniform particle view with hot spots in the front and top view.

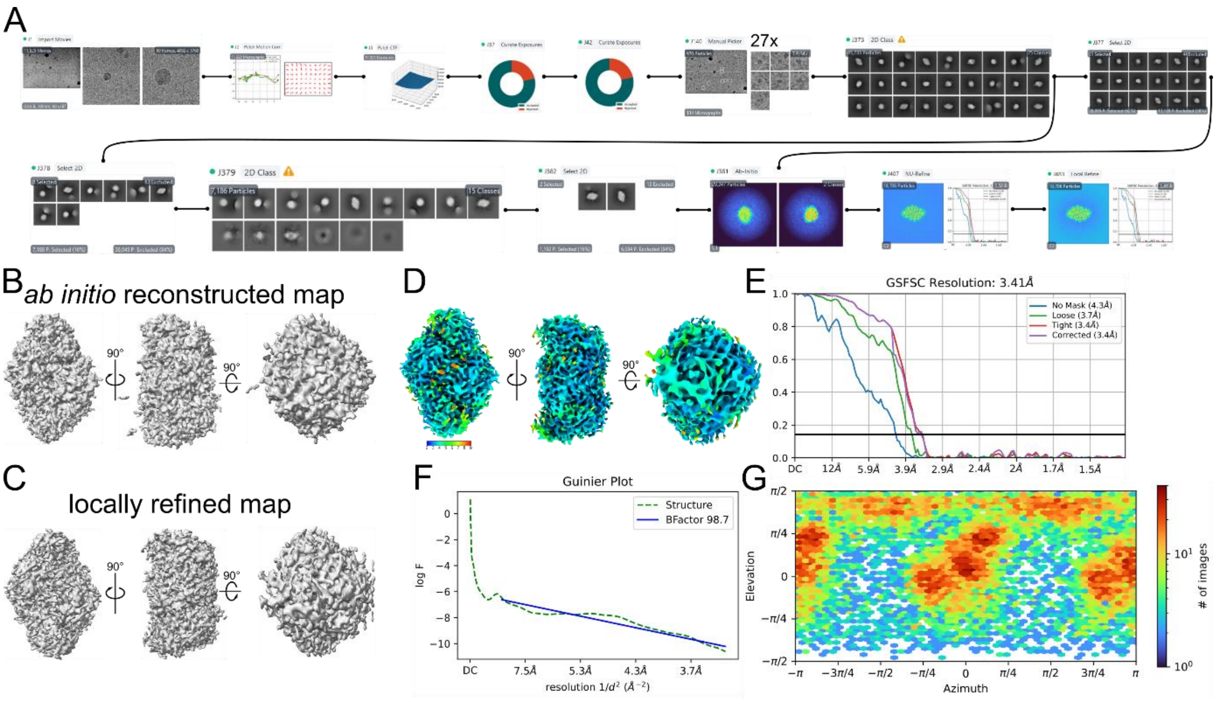

**Supplementary table 1 Constructs cloned, and tested for production, condensate formation, and catalysis.** The globular Rubisco (CbbM) is marked green, while all IDPs are marked grey. N or C suffixes indicate the terminal origin from the original protein. 05 indicates the first half of the respective IDP. ΔCtp indicates the deletion of the chloroplast targeting peptide from the original sequence.

| | |
|---|---|
| CbbM | DQSSRYVNLALKEEDLIAGGEHVLCAYIMKPKAGYGYVATAAHFAAESSTGTNVEVCTTDDFTRGVDALVYEVDEARELTKIAYPVALFDRNITDGKAMIASFLTLTMGNNQGMGDVEYAKMHDFYVPEAYRALFDGPSVNISALWKVLGRPEVDGGLVVGTIIKPKLGLRPKPFAEACHAFWLGGDFIKNDEPQGNQPFAPLRDTIALVADAMRRAQDETGEAKLFSANITADDPFEIIARGEYVLETFGENASHVALLVDGYVAGAAAITTARRRFPDNFLHYHRAGHGAVTSPQSKRGYTAFVHCKMARLQGASGIHTGTMGFGKMEGESSDRAIAYMLTQDEAQGPFYRQSWGGMKACTPIISGGMNALRMPGFFENLGNANVILTAGGGAFGHIDGPVAGARSLRQAWQAWRDGVPVLDYAREHKELARAFESFPGDADQIYPGWRKALGVEDTRSALPA |
| CbbM -Laf1N | DQSSRYVNLALKEEDLIAGGEHVLCAYIMKPKAGYGYVATAAHFAAESSTGTNVEVCTTDDFTRGVDALVYEVDEARELTKIAYPVALFDRNITDGKAMIASFLTLTMGNNQGMGDVEYAKMHDFYVPEAYRALFDGPSVNISALWKVLGRPEVDGGLVVGTIIKPKLGLRPKPFAEACHAFWLGGDFIKNDEPQGNQPFAPLRDTIALVADAMRRAQDETGEAKLFSANITADDPFEIIARGEYVLETFGENASHVALLVDGYVAGAAAITTARRRFPDNFLHYHRAGHGAVTSPQSKRGYTAFVHCKMARLQGASGIHTGTMGFGKMEGESSDRAIAYMLTQDEAQGPFYRQSWGGMKACTPIISGGMNALRMPGFFENLGNANVILTAGGGAFGHIDGPVAGARSLRQAWQAWRDGVPVLDYAREHKELARAFESFPGDADQIYPGWRKALGVEDTRSALPAESNQSNNGGSGNAALNRGGRYVPPHLRGGDGGAAAAASAGGDDRRGGAGGGGYRRGGGNSGGGGGGGYDRGYNDNRDDRDNRGGSGGYGRDRNYEDRGYNGGGGGGGNRGYNNNRGGGGGGYNRQDRGDGGSSNFSRGGYNNRDEGSDNRGSGRSYNNDRRDNGGDG |
| Laf1N- CbbM - Laf1N | ESNQSNNGGSGNAALNRGGRYVPPHLRGGDGGAAAAASAGGDDRRGGAGGGGYRRGGGNSGGGGGGGYDRGYNDNRDDRDNRGGSGGYGRDRNYEDRGYNGGGGGGGNRGYNNNRGGGGGGYNRQDRGDGGSSNFSRGGYNNRDEGSDNRGSGRSYNNDRRDNGGDGDQSSRYVNLALKEEDLIAGGEHVLCAYIMKPKAGYGYVATAAHFAAESSTGTNVEVCTTDDFTRGVDALVYEVDEARELTKIAYPVALFDRNITDGKAMIASFLTLTMGNNQGMGDVEYAKMHDFYVPEAYRALFDGPSVNISALWKVLGRPEVDGGLVVGTIIKPKLGLRPKPFAEACHAFWLGGDFIKNDEPQGNQPFAPLRDTIALVADAMRRAQDETGEAKLFSANITADDPFEIIARGEYVLETFGENASHVALLVDGYVAGAAAITTARRRFPDNFLHYHRAGHGAVTSPQSKRGYTAFVHCKMARLQGASGIHTGTMGFGKMEGESSDRAIAYMLTQDEAQGPFYRQSWGGMKACTPIISGGMNALRMPGFFENLGNANVILTAGGGAFGHIDGPVAGARSLRQAWQAWRDGVPVLDYAREHKELARAFESFPGDADQIYPGWRKALGVEDTRSALPAESNQSNNGGSGNAALNRGGRYVPPHLRGGDGGAAAAASAGGDDRRGGAGGGGYRRGGGNSGGGGGGGYDRGYNDNRDDRDNRGGSGGYGRDRNYEDRGYNGGGGGGGNRGYNNNRGGGGGGYNRQDRGDGGSSNFSRGGYNNRDEGSDNRGSGRSYNNDRRDNGGDG |
| CbbM - Laf1N05 | DQSSRYVNLALKEEDLIAGGEHVLCAYIMKPKAGYGYVATAAHFAAESSTGTNVEVCTTDDFTRGVDALVYEVDEARELTKIAYPVALFDRNITDGKAMIASFLTLTMGNNQGMGDVEYAKMHDFYVPEAYRALFDGPSVNISALWKVLGRPEVDGGLVVGTIIKPKLGLRPKPFAEACHAFWLGGDFIKNDEPQGNQPFAPLRDTIALVADAMRRAQDETGEAKLFSANITADDPFEIIARGEYVLETFGENASHVALLVDGYVAGAAAITTARRRFPDNFLHYHRAGHGAVTSPQSKRGYTAFVHCKMARLQGASGIHTGTMGFGKMEGESSDRAIAYMLTQDEAQGPFYRQSWGGMKACTPIISGGMNALRMPGFFENLGNANVILTAGGGAFGHIDGPVAGARSLRQAWQAWRDGVPVLDYAREHKELARAFESFPGDADQIYPGWRKALGVEDTRSALPAESNQSNNGGSGNAALNRGGRYVPPHLRGGDGGAAAAASAGGDDRRGGAGGGGYRRGGGNSGGGGGGGYDRGYNDNRDDRDNRGGSGGYGRDRNYEDRGYNGGGGGGGNRG |
| Laf1N05- CbbM - Laf1N05 | GSGGYGRDRNYEDRGYNGGGGGGGNRGYNNNRGGGGGGYNRQDRGDGGSSNFSRGGYNNRDEGSDNRGSGRSYNNDRRDNGGDGDQSSRYVNLALKEEDLIAGGEHVLCAYIMKPKAGYGYVATAAHFAAESSTGTNVEVCTTDDFTRGVDALVYEVDEARELTKIAYPVALFDRNITDGKAMIASFLTLTMGNNQGMGDVEYAKMHDFYVPEAYRALFDGPSVNISALWKVLGRPEVDGGLVVGTIIKPKLGLRPKPFAEACHAFWLGGDFIKNDEPQGNQPFAPLRDTIALVADAMRRAQDETGEAKLFSANITADDPFEIIARGEYVLETFGENASHVALLVDGYVAGAAAITTARRRFPDNFLHYHRAGHGAVTSPQSKRGYTAFVHCKMARLQGASGIHTGTMGFGKMEGESSDRAIAYMLTQDEAQGPFYRQSWGGMKACTPIISGGMNALRMPGFFENLGNANVILTAGGGAFGHIDGPVAGARSLRQAWQAWRDGVPVLDYAREHKELARAFESFPGDADQIYPGWRKALGVEDTRSALPAESNQSNNGGSGNAALNRGGRYVPPHLRGGDGGAAAAASAGGDDRRGGAGGGGYRRGGGNSGGGGGGGYDRGYNDNRDDRDNRG |
| CbbM -Ddx4N | DQSSRYVNLALKEEDLIAGGEHVLCAYIMKPKAGYGYVATAAHFAAESSTGTNVEVCTTDDFTRGVDALVYEVDEARELTKIAYPVALFDRNITDGKAMIASFLTLTMGNNQGMGDVEYAKMHDFYVPEAYRALFDGPSVNISALWKVLGRPEVDGGLVVGTIIKPKLGLRPKPFAEACHAFWLGGDFIKNDEPQGNQPFAPLRDTIALVADAMRRAQDETGEAKLFSANITADDPFEIIARGEYVLETFGENASHVALLVDGYVAGAAAITTARRRFPDNFLHYHRAGHGAVTSPQSKRGYTAFVHCKMARLQGASGIHTGTMGFGKMEGESSDRAIAYMLTQDEAQGPFYRQSWGGMKACTPIISGGMNALRMPGFFENLGNANVILTAGGGAFGHIDGPVAGARSLRQAWQAWRDGVPVLDYAREHKELARAFESFPGDADQIYPGWRKALGVEDTRSALPAGDEDWEAEINPHMSSYVPIFEKDRYSGENGDNFNRTPASSSEMDDGPSRRDHFMKSGFASGRNFGNRDAGECNKRDNTSTMGGFGVGKSFGNRGFSNSRFEDGDSSGFWRESSNDCEDNPTRNRGFSKRGGYRDGNNSEASGPYRRGGRGSFRGCRGGFGLGSPNNDLDPDECMQRTGGLFGSRRPVLSGTGNGDTSQSRSGSGSERGGYKGLNEEVITGSGKNSWKSEAEGGES |
| Ddx4N- CbbM -Ddx4N | GDEDWEAEINPHMSSYVPIFEKDRYSGENGDNFNRTPASSSEMDDGPSRRDHFMKSGFASGRNFGNRDAGECNKRDNTSTMGGFGVGKSFGNRGFSNSRFEDGDSSGFWRESSNDCEDNPTRNRGFSKRGGYRDGNNSEASGPYRRGGRGSFRGCRGGFGLGSPNNDLDPDECMQRTGGLFGSRRPVLSGTGNGDTSQSRSGSGSERGGYKGLNEEVITGSGKNSWKSEAEGGESDQSSRYVNLALKEEDLIAGGEHVLCAYIMKPKAGYGYVATAAHFAAESSTGTNVEVCTTDDFTRGVDALVYEVDEARELTKIAYPVALFDRNITDGKAMIASFLTLTMGNNQGMGDVEYAKMHDFYVPEAYRALFDGPSVNISALWKVLGRPEVDGGLVVGTIIKPKLGLRPKPFAEACHAFWLGGDFIKNDEPQGNQPFAPLRDTIALVADAMRRAQDETGEAKLFSANITADDPFEIIARGEYVLETFGENASHVALLVDGYVAGAAAITTARRRFPDNFLHYHRAGHGAVTSPQSKRGYTAFVHCKMARLQGASGIHTGTMGFGKMEGESSDRAIAYMLTQDEAQGPFYRQSWGGMKACTPIISGGMNALRMPGFFENLGNANVILTAGGGAFGHIDGPVAGARSLRQAWQAWRDGVPVLDYAREHKELARAFESFPGDADQIYPGWRKALGVEDTRSALPAGDEDWEAEINPHMSSYVPIFEKDRYSGENGDNFNRTPASSSEMDDGPSRRDHFMKSGFASGRNFGNRDAGECNKRDNTSTMGGFGVGKSFGNRGFSNSRFEDGDSSGFWRESSNDCEDNPTRNRGFSKRGGYRDGNNSEASGPYRRGGRGSFRGCRGGFGLGSPNNDLDPDECMQRTGGLFGSRRPVLSGTGNGDTSQSRSGSGSERGGYKGLNEEVITGSGKNSWKSEAEGGES |
| CbbM -UBQLN2C | DQSSRYVNLALKEEDLIAGGEHVLCAYIMKPKAGYGYVATAAHFAAESSTGTNVEVCTTDDFTRGVDALVYEVDEARELTKIAYPVALFDRNITDGKAMIASFLTLTMGNNQGMGDVEYAKMHDFYVPEAYRALFDGPSVNISALWKVLGRPEVDGGLVVGTIIKPKLGLRPKPFAEACHAFWLGGDFIKNDEPQGNQPFAPLRDTIALVADAMRRAQDETGEAKLFSANITADDPFEIIARGEYVLETFGENASHVALLVDGYVAGAAAITTARRRFPDNFLHYHRAGHGAVTSPQSKRGYTAFVHCKMARLQGASGIHTGTMGFGKMEGESSDRAIAYMLTQDEAQGPFYRQSWGGMKACTPIISGGMNALRMPGFFENLGNANVILTAGGGAFGHIDGPVAGARSLRQAWQAWRDGVPVLDYAREHKELARAFESFPGDADQIYPGWRKALGVEDTRSALPAAMQALMQIQQGLQTLATEAPGLIPSFTPGVGVGVLGTAIGPVGPVTPIGPIGPIVPFTPIGPIGPIGPTGPAAPPGSTGSGGPTGPTVSSAAPSETTSPTSESGPNQQFIQQMVQALAGANAPQLPNPEVRFQQQLEQLNAMGFLNREANLQALIATGGDINAAIERLLGSQPS |
| CbbM -UBQLN2C05 | DQSSRYVNLALKEEDLIAGGEHVLCAYIMKPKAGYGYVATAAHFAAESSTGTNVEVCTTDDFTRGVDALVYEVDEARELTKIAYPVALFDRNITDGKAMIASFLTLTMGNNQGMGDVEYAKMHDFYVPEAYRALFDGPSVNISALWKVLGRPEVDGGLVVGTIIKPKLGLRPKPFAEACHAFWLGGDFIKNDEPQGNQPFAPLRDTIALVADAMRRAQDETGEAKLFSANITADDPFEIIARGEYVLETFGENASHVALLVDGYVAGAAAITTARRRFPDNFLHYHRAGHGAVTSPQSKRGYTAFVHCKMARLQGASGIHTGTMGFGKMEGESSDRAIAYMLTQDEAQGPFYRQSWGGMKACTPIISGGMNALRMPGFFENLGNANVILTAGGGAFGHIDGPVAGARSLRQAWQAWRDGVPVLDYAREHKELARAFESFPGDADQIYPGWRKALGVEDTRSALPAAMQALMQIQQGLQTLATEAPGLIPSFTPGVGVGVLGTAIGPVGPVTPIGPIGPIVPFTPIGPIGPIGPTGPAAPPGSTGSGGPTGP |
| UBQLN2C- CbbM - UBQLN2C | AMQALMQIQQGLQTLATEAPGLIPSFTPGVGVGVLGTAIGPVGPVTPIGPIGPIVPFTPIGPIGPIGPTGPAAPPGSTGSGGPTGPTVSSAAPSETTSPTSESGPNQQFIQQMVQALAGANAPQLPNPEVRFQQQLEQLNAMGFLNREANLQALIATGGDINAAIERLLGSQPSDQSSRYVNLALKEEDL |

| | |
|---|---|
| | IAGGEHVLCAYIMKPKAGYGYVATAAHFAAESSTGTNVEVCTTDDFTRGVDALVYEVDEARELTKIAYPVALFDRNITDGKAMIASFLTLTMGNNQGMGDVEYAKMHDFYVPEAYRALFDGPSVNISALWKVLGRPEVDGGLVVGTIIKPKLGLRPKPFAEACHAFWLGGDFIKNDEPQGNQPFAPLRDTIALVADAMRRAQDETGEAKLFSANITADDPFEIIARGEYVLETFGENASHVALLVDGYVAGAAAITTARRRFPDNFLHYHRAGHGAVTSPQSKRGYTAFVHCKMARLQGASGIHTGTMGFGKMEGESSDRAIAYMLTQDEAQGPFYRQSWGGMKACTPIISGGMNALRMPGFFENLGNANVILTAGGGAFGHIDGPVAGARSLRQAWQAWRDGVPVLDYAREHKELARAFESFPGDADQIYPGWRKALGVEDTRSALPAAMQALMQIQQGLQTLATEAPGLIPSFTPGVGVGVLGTAIGPVGPVTPIGPIGPIVPFTPIGPIGPIGPTGPAAPPGSTGSGGPTGPTVSSAAPSETTSPTSESGPNQQFIQQMVQALAGANAPQLPNPEVRFQQQLEQLNAMGFLNREANLQALIATGGDINAAIERLLGSQPS |
| CbbM-GATA3N | DQSSRYVNLALKEEDLIAGGEHVLCAYIMKPKAGYGYVATAAHFAAESSTGTNVEVCTTDDFTRGVDALVYEVDEARELTKIAYPVALFDRNITDGKAMIASFLTLTMGNNQGMGDVEYAKMHDFYVPEAYRALFDGPSVNISALWKVLGRPEVDGGLVVGTIIKPKLGLRPKPFAEACHAFWLGGDFIKNDEPQGNQPFAPLRDTIALVADAMRRAQDETGEAKLFSANITADDPFEIIARGEYVLETFGENASHVALLVDGYVAGAAAITTARRRFPDNFLHYHRAGHGAVTSPQSKRGYTAFVHCKMARLQGASGIHTGTMGFGKMEGESSDRAIAYMLTQDEAQGPFYRQSWGGMKACTPIISGGMNALRMPGFFENLGNANVILTAGGGAFGHIDGPVAGARSLRQAWQAWRDGVPVLDYAREHKELARAFESFPGDADQIYPGWRKALGVEDTRSALPAVTADQPRWVSHHHPAVLNGQHPDTHHPGLSHSYMDAAQYPLPEEVDVLFNIDGQGNHVPPYYGNSVRATVQRYPPTHHGSQVCRPPLLHGSLPWLDGGKALGSHHTASPWNLSPFSKTSIHHGSPGPLSVYPPASSSSLSGGHASPHLFTFPPTPPKDVSPDPSLSTPGSAGSARQDEKECLKYQVPLPDSMKLESSHSRGSMTALGGASSSTHHPITTYPPYVPEYSSGLFPPSSLLGGSPTGFGCK |
| CbbM-GATA3N05 | DQSSRYVNLALKEEDLIAGGEHVLCAYIMKPKAGYGYVATAAHFAAESSTGTNVEVCTTDDFTRGVDALVYEVDEARELTKIAYPVALFDRNITDGKAMIASFLTLTMGNNQGMGDVEYAKMHDFYVPEAYRALFDGPSVNISALWKVLGRPEVDGGLVVGTIIKPKLGLRPKPFAEACHAFWLGGDFIKNDEPQGNQPFAPLRDTIALVADAMRRAQDETGEAKLFSANITADDPFEIIARGEYVLETFGENASHVALLVDGYVAGAAAITTARRRFPDNFLHYHRAGHGAVTSPQSKRGYTAFVHCKMARLQGASGIHTGTMGFGKMEGESSDRAIAYMLTQDEAQGPFYRQSWGGMKACTPIISGGMNALRMPGFFENLGNANVILTAGGGAFGHIDGPVAGARSLRQAWQAWRDGVPVLDYAREHKELARAFESFPGDADQIYPGWRKALGVEDTRSALPAVTADQPRWVSHHHPAVLNGQHPDTHHPGLSHSYMDAAQYPLPEEVDVLFNIDGQGNHVPPYYGNSVRATVQRYPPTHHGSQVCRPPLLHGSLPWLDGGKALGSHHTASPWNLSPFSKTSIHHGS |
| GATA3N-CbbM-GATA3N | VTADQPRWVSHHHPAVLNGQHPDTHHPGLSHSYMDAAQYPLPEEVDVLFNIDGQGNHVPPYYGNSVRATVQRYPPTHHGSQVCRPPLLHGSLPWLDGGKALGSHHTASPWNLSPFSKTSIHHGSPGPLSVYPPASSSSLSGGHASPHLFTFPPTPPKDVSPDPSLSTPGSAGSARQDEKECLKYQVPLPDSMKLESSHSRGSMTALGGASSSTHHPITTYPPYVPEYSSGLFPPSSLLGGSPTGFGCKDQSSRYVNLALKEEDLIAGGEHVLCAYIMKPKAGYGYVATAAHFAAESSTGTNVEVCTTDDFTRGVDALVYEVDEARELTKIAYPVALFDRNITDGKAMIASFLTLTMGNNQGMGDVEYAKMHDFYVPEAYRALFDGPSVNISALWKVLGRPEVDGGLVVGTIIKPKLGLRPKPFAEACHAFWLGGDFIKNDEPQGNQPFAPLRDTIALVADAMRRAQDETGEAKLFSANITADDPFEIIARGEYVLETFGENASHVALLVDGYVAGAAAITTARRRFPDNFLHYHRAGHGAVTSPQSKRGYTAFVHCKMARLQGASGIHTGTMGFGKMEGESSDRAIAYMLTQDEAQGPFYRQSWGGMKACTPIISGGMNALRMPGFFENLGNANVILTAGGGAFGHIDGPVAGARSLRQAWQAWRDGVPVLDYAREHKELARAFESFPGDADQIYPGWRKALGVEDTRSALPAVTADQPRWVSHHHPAVLNGQHPDTHHPGLSHSYMDAAQYPLPEEVDVLFNIDGQGNHVPPYYGNSVRATVQRYPPTHHGSQVCRPPLLHGSLPWLDGGKALGSHHTASPWNLSPFSKTSIHHGSPGPLSVYPPASSSSLSGGHASPHLFTFPPTPPKDVSPDPSLSTPGSAGSARQDEKECLKYQVPLPDSMKLESSHSRGSMTALGGASSSTHHPITTYPPYVPEYSSGLFPPSSLLGGSPTGFGCK |
| CbbM-AKAP95N | DQSSRYVNLALKEEDLIAGGEHVLCAYIMKPKAGYGYVATAAHFAAESSTGTNVEVCTTDDFTRGVDALVYEVDEARELTKIAYPVALFDRNITDGKAMIASFLTLTMGNNQGMGDVEYAKMHDFYVPEAYRALFDGPSVNISALWKVLGRPEVDGGLVVGTIIKPKLGLRPKPFAEACHAFWLGGDFIKNDEPQGNQPFAPLRDTIALVADAMRRAQDETGEAKLFSANITADDPFEIIARGEYVLETFGENASHVALLVDGYVAGAAAITTARRRFPDNFLHYHRAGHGAVTSPQSKRGYTAFVHCKMARLQGASGIHTGTMGFGKMEGESSDRAIAYMLTQDEAQGPFYRQSWGGMKACTPIISGGMNALRMPGFFENLGNANVILTAGGGAFGHIDGPVAGARSLRQAWQAWRDGVPVLDYAREHKELARAFESFPGDADQIYPGWRKALGVEDTRSALPADMMSKEGGRGGSGGGGEGIQDRESSFRFQPFESYDSRPCLPEHNPYRPSYSYDYEFDLGSDRNGSFGGQYSECRDPARERGSLDGFMRGRGQGRFQDRSNPGTFMRSDPF |
| CbbM-AKAP95N05 | DQSSRYVNLALKEEDLIAGGEHVLCAYIMKPKAGYGYVATAAHFAAESSTGTNVEVCTTDDFTRGVDALVYEVDEARELTKIAYPVALFDRNITDGKAMIASFLTLTMGNNQGMGDVEYAKMHDFYVPEAYRALFDGPSVNISALWKVLGRPEVDGGLVVGTIIKPKLGLRPKPFAEACHAFWLGGDFIKNDEPQGNQPFAPLRDTIALVADAMRRAQDETGEAKLFSANITADDPFEIIARGEYVLETFGENASHVALLVDGYVAGAAAITTARRRFPDNFLHYHRAGHGAVTSPQSKRGYTAFVHCKMARLQGASGIHTGTMGFGKMEGESSDRAIAYMLTQDEAQGPFYRQSWGGMKACTPIISGGMNALRMPGFFENLGNANVILTAGGGAFGHIDGPVAGARSLRQAWQAWRDGVPVLDYAREHKELARAFESFPGDADQIYPGWRKALGVEDTRSALPADMMSKEGGRGGSGGGGEGIQDRESSFRFQPFESYDSRPCLPEHNPYRPSYSYDYE |
| AKAP95N-CbbM-AKAP95N | DMMSKEGGRGGSGGGGEGIQDRESSFRFQPFESYDSRPCLPEHNPYRPSYSYDYEFDLGSDRNGSFGGQYSECRDPARERGSLDGFMRGRGQGRFQDRSNPGTFMRSDPFDQSSRYVNLALKEEDLIAGGEHVLCAYIMKPKAGYGYVATAAHFAAESSTGTNVEVCTTDDFTRGVDALVYEVDEARELTKIAYPVALFDRNITDGKAMIASFLTLTMGNNQGMGDVEYAKMHDFYVPEAYRALFDGPSVNISALWKVLGRPEVDGGLVVGTIIKPKLGLRPKPFAEACHAFWLGGDFIKNDEPQGNQPFAPLRDTIALVADAMRRAQDETGEAKLFSANITADDPFEIIARGEYVLETFGENASHVALLVDGYVAGAAAITTARRRFPDNFLHYHRAGHGAVTSPQSKRGYTAFVHCKMARLQGASGIHTGTMGFGKMEGESSDRAIAYMLTQDEAQGPFYRQSWGGMKACTPIISGGMNALRMPGFFENLGNANVILTAGGGAFGHIDGPVAGARSLRQAWQAWRDGVPVLDYAREHKELARAFESFPGDADQIYPGWRKALGVEDTRSALPADMMSKEGGRGGSGGGGEGIQDRESSFRFQPFESYDSRPCLPEHNPYRPSYSYDYEFDLGSDRNGSFGGQYSECRDPARERGSLDGFMRGRGQGRFQDRSNPGTFMRSDPF |
| CbbM-FXR1C | DQSSRYVNLALKEEDLIAGGEHVLCAYIMKPKAGYGYVATAAHFAAESSTGTNVEVCTTDDFTRGVDALVYEVDEARELTKIAYPVALFDRNITDGKAMIASFLTLTMGNNQGMGDVEYAKMHDFYVPEAYRALFDGPSVNISALWKVLGRPEVDGGLVVGTIIKPKLGLRPKPFAEACHAFWLGGDFIKNDEPQGNQPFAPLRDTIALVADAMRRAQDETGEAKLFSANITADDPFEIIARGEYVLETFGENASHVALLVDGYVAGAAAITTARRRFPDNFLHYHRAGHGAVTSPQSKRGYTAFVHCKMARLQGASGIHTGTMGFGKMEGESSDRAIAYMLTQDEAQGPFYRQSWGGMKACTPIISGGMNALRMPGFFENLGNANVILTAGGGAFGHIDGPVAGARSLRQAWQAWRDGVPVLDYAREHKELARAFESFPGDADQIYPGWRKALGVEDTRSALPAQLRQIGMGFRPSSSRGTEKEKGYATDESTASSVRGSRSYSGRGRGRRGPNYTSGYGTNSELSNPSETESERKEELSDWSLAGEDERESRQQRDSRRRPGGRGRSGSAGRGRGGSRGGKSSISSVLKDPDSNPYSLLDNTESDQTADTDASESHHNTNRRRRSRRRRTDEDSSLMDGMTESDNASVNENGLDDSEQKPQRRNRSRRRRFRGQAEDRQPVTVADYISRAESQSRQRNLPKEPLAKGKKEKVKDVIEEHGPSEKVINGPRAASADKALKPQTTERNKASCQDGSKQEAILNGVS |
| FXR1C-CbbM-FXR1C | QLRQIGMGFRPSSSRGTEKEKGYATDESTASSVRGSRSYSGRGRGRRGPNYTSGYGTNSELSNPSETESERKEELSDWSLAGEDERESRQQRDSRRRPGGRGRSGSAGRGRGGSRGGKSSISSVLKDPDSNPYSLLDNTESDQTADTDASESHHNTNRRRRSRRRRTDEDSSLMDGMTESDNASVNENGLDDSEQKPQRRNRSRRRRFRGQAEDRQPVTVADYISRAESQSRQRNLPKEPLAKGKKEKVKDVIEEHGPSEKVINGPRAASADKALKPQTTERNKASCQDGSKQEAILNGVSDQSSRYVNLALKEEDLIAGGEHVLCAYIMKPKAGYGYVATAAHFAAESSTGTNVEVCTTDDFTRGVDALVYEVDEARELTKIAYPVALFDRNITDGKAMIASFLTLTMGNNQGMGDVEYAKMHDFYVPEAYRALFDGPSVNISALWKVLGRPEVDGGLVVGTIIKPKLGLRPKPFAEACHAFWLGGDFIKNDEPQGNQPFAPLRDTIALVADAMRRAQDETGEAKLFSANITADDPFEIIARGEYVLETFGENASHVALLVDGYVAGAAAITTARRRFPDNFLHYHRAGHGAVTSPQSKRGYTAFVHCKMARLQGASGIHTGTMGFGKMEGESSDRAIAYMLTQDEAQGPFYRQSWGGMKACTPIISGGMNALRMPGFFENLGNANVILTAGGGAFGHIDGPVAGARSLRQAWQAWRDGVPVLDYAREHKELARAFESFPGDADQIYPGWRKALGVEDTRSALPAQLRQIGMGFRPSSSRGTEKEKGYATDESTASSVRGSRSYSGRGRGRRGPNYTSGYGTNSELSNPSETESERKEELSDWSLAGEDERESRQQRDSRRRPGGRGRSGSAGRGRGGSRGGKSSISSVLKDPDSNPYSLLDNTESDQTADTDASESHHNTNRRRRSRRRRTDEDSSLMDGMTESDNASVNENGLDDSEQKPQRRNRSRRRRFRGQAEDRQPVTVADYISRAESQSRQRNLPKEPLAKGKKEKVKDVIEEHGPSEKVINGPRAASADKALKPQTTERNKASCQDGSKQEAILNGVS |

| | |
|---|---|
| CbbM-LEM2N | DQSSRYVNLALKEEDLIAGGEHVLCAYIMKPKAGYGYVATAAHFAAESSTGTNVEVCTTDDFTRGVDALVYEVDEARELTKIAYPVALFDRNITDG<br>KAMIASFLTLTMGNNQGMGDVEYAKMHDFYVPEAYRALFDGPSVNISALWKVLGRPEVDGGLVVGTIIKPKLGLRPKPFAEACHAFWLGGDFIK<br>NDEPQGNQPFAPLRDTIALVADAMRRAQDETGEAKLFSANITADDPFEIIARGEYVLETFGENASHVALLVDGYVAGAAAITTARRRFPDNFLHY<br>HRAGHGAVTSPQSKRGYTAFVHCKMARLQGASGIHTGTMGFGKMEGESSDRAIAYMLTQDEAQGPFYRQSWGGMKACTPIISGGMNALRM<br>PGFFENLGNANVILTAGGGAFGHIDGPVAGARSLRQAWQAWRDGVPVLDYAREHKELARAFESFPGDADQIYPGWRKALGVEDTRSALPARL<br>RDEERLREEARPRGEERLREEARLREDAPLRARPAAASPRAEPWLSQPASGSAYATPGAYGDIRPSAASWVGSRGLAYPARPAQLRRRASVRGSS<br>EEDEDARTPDRATQGPGLAARRWWAASPAPARLPSSLLGPDPRPGLRATRAGPAGAARARPEV |
| LEM2N-CbbM-LEM2N | RLRDEERLREEARPRGEERLREEARLREDAPLRARPAAASPRAEPWLSQPASGSAYATPGAYGDIRPSAASWVGSRGLAYPARPAQLRRRASVRG<br>SSEEDEDARTPDRATQGPGLAARRWWAASPAPARLPSSLLGPDPRPGLRATRAGPAGAARARPEVDQSSRYVNLALKEEDLIAGGEHVLCAYIM<br>KPKAGYGYVATAAHFAAESSTGTNVEVCTTDDFTRGVDALVYEVDEARELTKIAYPVALFDRNITDGKAMIASFLTLTMGNNQGMGDVEYAKMH<br>DFYVPEAYRALFDGPSVNISALWKVLGRPEVDGGLVVGTIIKPKLGLRPKPFAEACHAFWLGGDFIKNDEPQGNQPFAPLRDTIALVADAMRRAQ<br>DETGEAKLFSANITADDPFEIIARGEYVLETFGENASHVALLVDGYVAGAAAITTARRRFPDNFLHYHRAGHGAVTSPQSKRGYTAFVHCKMARL<br>QGASGIHTGTMGFGKMEGESSDRAIAYMLTQDEAQGPFYRQSWGGMKACTPIISGGMNALRMPGFFENLGNANVILTAGGGAFGHIDGPVA<br>GARSLRQAWQAWRDGVPVLDYAREHKELARAFESFPGDADQIYPGWRKALGVEDTRSALPARLRDEERLREEARPRGEERLREEARLREDAPL<br>RARPAAASPRAEPWLSQPASGSAYATPGAYGDIRPSAASWVGSRGLAYPARPAQLRRRASVRGSSEEDEDARTPDRATQGPGLAARRWWAASP<br>APARLPSSLLGPDPRPGLRATRAGPAGAARARPEV |
| CbbM-xCPEB4N | DQSSRYVNLALKEEDLIAGGEHVLCAYIMKPKAGYGYVATAAHFAAESSTGTNVEVCTTDDFTRGVDALVYEVDEARELTKIAYPVALFDRNITDG<br>KAMIASFLTLTMGNNQGMGDVEYAKMHDFYVPEAYRALFDGPSVNISALWKVLGRPEVDGGLVVGTIIKPKLGLRPKPFAEACHAFWLGGDFIK<br>NDEPQGNQPFAPLRDTIALVADAMRRAQDETGEAKLFSANITADDPFEIIARGEYVLETFGENASHVALLVDGYVAGAAAITTARRRFPDNFLHY<br>HRAGHGAVTSPQSKRGYTAFVHCKMARLQGASGIHTGTMGFGKMEGESSDRAIAYMLTQDEAQGPFYRQSWGGMKACTPIISGGMNALRM<br>PGFFENLGNANVILTAGGGAFGHIDGPVAGARSLRQAWQAWRDGVPVLDYAREHKELARAFESFPGDADQIYPGWRKALGVEDTRSALPAGD<br>YGFGVLVQSNTGNKSAFPVRFHPHLQPPHHHQNATPSPAAFINNNTAANGSSAGSAWLFPAPATHNIQDEILGSEKAKSQQQEQQDPLEKQQL<br>SPSPGQEAGILPETEKAKSEENQGDNSSENGNGKEKIRIESPVLTGFDYQEATGLGTSTQPLTSSASSLTGFSNWSAAIAPSSSTIINEDASFFHQGG<br>VPAASANN |
| CbbM-xCPEB4N05 | DQSSRYVNLALKEEDLIAGGEHVLCAYIMKPKAGYGYVATAAHFAAESSTGTNVEVCTTDDFTRGVDALVYEVDEARELTKIAYPVALFDRNITDG<br>KAMIASFLTLTMGNNQGMGDVEYAKMHDFYVPEAYRALFDGPSVNISALWKVLGRPEVDGGLVVGTIIKPKLGLRPKPFAEACHAFWLGGDFIK<br>NDEPQGNQPFAPLRDTIALVADAMRRAQDETGEAKLFSANITADDPFEIIARGEYVLETFGENASHVALLVDGYVAGAAAITTARRRFPDNFLHY<br>HRAGHGAVTSPQSKRGYTAFVHCKMARLQGASGIHTGTMGFGKMEGESSDRAIAYMLTQDEAQGPFYRQSWGGMKACTPIISGGMNALRM<br>PGFFENLGNANVILTAGGGAFGHIDGPVAGARSLRQAWQAWRDGVPVLDYAREHKELARAFESFPGDADQIYPGWRKALGVEDTRSALPAGD<br>YGFGVLVQSNTGNKSAFPVRFHPHLQPPHHHQNATPSPAAFINNNTAANGSSAGSAWLFPAPATHNIQDEILGSEKAKSQQQEQQDPLEKQQL<br>SPSPGQ |
| xCPEB4N-CbbM-xCPEB4N | GDYGFGVLVQSNTGNKSAFPVRFHPHLQPPHHHQNATPSPAAFINNNTAANGSSAGSAWLFPAPATHNIQDEILGSEKAKSQQQEQQDPLEKQ<br>QLSPSPGQEAGILPETEKAKSEENQGDNSSENGNGKEKIRIESPVLTGFDYQEATGLGTSTQPLTSSASSLTGFSNWSAAIAPSSSTIINEDASFFHQ<br>GGVPAASANNDQSSRYVNLALKEEDLIAGGEHVLCAYIMKPKAGYGYVATAAHFAAESSTGTNVEVCTTDDFTRGVDALVYEVDEARELTKIAYP<br>VALFDRNITDGKAMIASFLTLTMGNNQGMGDVEYAKMHDFYVPEAYRALFDGPSVNISALWKVLGRPEVDGGLVVGTIIKPKLGLRPKPFAEAC<br>HAFWLGGDFIKNDEPQGNQPFAPLRDTIALVADAMRRAQDETGEAKLFSANITADDPFEIIARGEYVLETFGENASHVALLVDGYVAGAAAITTA<br>RRRFPDNFLHYHRAGHGAVTSPQSKRGYTAFVHCKMARLQGASGIHTGTMGFGKMEGESSDRAIAYMLTQDEAQGPFYRQSWGGMKACTPI<br>ISGGMNALRMPGFFENLGNANVILTAGGGAFGHIDGPVAGARSLRQAWQAWRDGVPVLDYAREHKELARAFESFPGDADQIYPGWRKALGV<br>EDTRSALPAGDYGFGVLVQSNTGNKSAFPVRFHPHLQPPHHHQNATPSPAAFINNNTAANGSSAGSAWLFPAPATHNIQDEILGSEKAKSQQQE<br>QQDPLEKQQLSPSPGQEAGILPETEKAKSEENQGDNSSENGNGKEKIRIESPVLTGFDYQEATGLGTSTQPLTSSASSLTGFSNWSAAIAPSSSTIIN<br>EDASFFHQGGVPAASANN |
| CbbM-U2AF65N | DQSSRYVNLALKEEDLIAGGEHVLCAYIMKPKAGYGYVATAAHFAAESSTGTNVEVCTTDDFTRGVDALVYEVDEARELTKIAYPVALFDRNITDG<br>KAMIASFLTLTMGNNQGMGDVEYAKMHDFYVPEAYRALFDGPSVNISALWKVLGRPEVDGGLVVGTIIKPKLGLRPKPFAEACHAFWLGGDFIK<br>NDEPQGNQPFAPLRDTIALVADAMRRAQDETGEAKLFSANITADDPFEIIARGEYVLETFGENASHVALLVDGYVAGAAAITTARRRFPDNFLHY<br>HRAGHGAVTSPQSKRGYTAFVHCKMARLQGASGIHTGTMGFGKMEGESSDRAIAYMLTQDEAQGPFYRQSWGGMKACTPIISGGMNALRM<br>PGFFENLGNANVILTAGGGAFGHIDGPVAGARSLRQAWQAWRDGVPVLDYAREHKELARAFESFPGDADQIYPGWRKALGVEDTRSALPASD<br>FDEFERQLNENKQERDKENRHRKRSHSRSRSRDRKRRSRSRDRRNRDQRSASRDRRRRSKPLTRGAKEEHGGLIRSPRHEKKKKVRKYWDVPPP<br>GFEHITPMQYKAMQAAGQIPATALLPTMTPDGLAVTPTPVPVVGSQMTRQAR |
| CbbM-U2AF65N05 | DQSSRYVNLALKEEDLIAGGEHVLCAYIMKPKAGYGYVATAAHFAAESSTGTNVEVCTTDDFTRGVDALVYEVDEARELTKIAYPVALFDRNITDG<br>KAMIASFLTLTMGNNQGMGDVEYAKMHDFYVPEAYRALFDGPSVNISALWKVLGRPEVDGGLVVGTIIKPKLGLRPKPFAEACHAFWLGGDFIK<br>NDEPQGNQPFAPLRDTIALVADAMRRAQDETGEAKLFSANITADDPFEIIARGEYVLETFGENASHVALLVDGYVAGAAAITTARRRFPDNFLHY<br>HRAGHGAVTSPQSKRGYTAFVHCKMARLQGASGIHTGTMGFGKMEGESSDRAIAYMLTQDEAQGPFYRQSWGGMKACTPIISGGMNALRM<br>PGFFENLGNANVILTAGGGAFGHIDGPVAGARSLRQAWQAWRDGVPVLDYAREHKELARAFESFPGDADQIYPGWRKALGVEDTRSALPSDF<br>DEFERQLNENKQERDKENRHRKRSHSRSRSRDRKRRSRSRDRRNRDQRSASRDRRRRSKPLTRGAKEEHGG |
| U2AF65N-CbbM-<br>U2AF65N | SDFDEFERQLNENKQERDKENRHRKRSHSRSRSRDRKRRSRSRDRRNRDQRSASRDRRRRSKPLTRGAKEEHGGLIRSPRHEKKKKVRKYWDVP<br>PPGFEHITPMQYKAMQAAGQIPATALLPTMTPDGLAVTPTPVPVVGSQMTRQARDQSSRYVNLALKEEDLIAGGEHVLCAYIMKPKAGYGYVA<br>TAAHFAAESSTGTNVEVCTTDDFTRGVDALVYEVDEARELTKIAYPVALFDRNITDGKAMIASFLTLTMGNNQGMGDVEYAKMHDFYVPEAYRA<br>LFDGPSVNISALWKVLGRPEVDGGLVVGTIIKPKLGLRPKPFAEACHAFWLGGDFIKNDEPQGNQPFAPLRDTIALVADAMRRAQDETGEAKLFS<br>ANITADDPFEIIARGEYVLETFGENASHVALLVDGYVAGAAAITTARRRFPDNFLHYHRAGHGAVTSPQSKRGYTAFVHCKMARLQGASGIHTGT<br>MGFGKMEGESSDRAIAYMLTQDEAQGPFYRQSWGGMKACTPIISGGMNALRMPGFFENLGNANVILTAGGGAFGHIDGPVAGARSLRQAW<br>QAWRDGVPVLDYAREHKELARAFESFPGDADQIYPGWRKALGVEDTRSALPASDFDEFERQLNENKQERDKENRHRKRSHSRSRSRDRKRRSR<br>SRDRRNRDQRSASRDRRRRSKPLTRGAKEEHGGLIRSPRHEKKKKVRKYWDVPPPGFEHITPMQYKAMQAAGQIPATALLPTMTPDGLAVTPT<br>PVPVVGSQMTRQAR |
| CbbM-EPYC1AGΔCtp | DQSSRYVNLALKEEDLIAGGEHVLCAYIMKPKAGYGYVATAAHFAAESSTGTNVEVCTTDDFTRGVDALVYEVDEARELTKIAYPVALFDRNITDG<br>KAMIASFLTLTMGNNQGMGDVEYAKMHDFYVPEAYRALFDGPSVNISALWKVLGRPEVDGGLVVGTIIKPKLGLRPKPFAEACHAFWLGGDFIK<br>NDEPQGNQPFAPLRDTIALVADAMRRAQDETGEAKLFSANITADDPFEIIARGEYVLETFGENASHVALLVDGYVAGAAAITTARRRFPDNFLHY<br>HRAGHGAVTSPQSKRGYTAFVHCKMARLQGASGIHTGTMGFGKMEGESSDRAIAYMLTQDEAQGPFYRQSWGGMKACTPIISGGMNALRM<br>PGFFENLGNANVILTAGGGAFGHIDGPVAGARSLRQAWQAWRDGVPVLDYAREHKELARAFESFPGDADQIYPGWRKALGVEDTRSALPAAR<br>GSWRDSSSVTTKVPSRSSSSANRVSPTRSVLPANWRQELESLRNGNGSAPSAPTPAAPARSSSANWRTESSSSAPKPASSAPARPSSSPRRTVTP<br>NRSSLPPNWKQELESLRNGNGNGSSSSYSSYSAPAPARSSSASWRTEAAPAAPAAAARPSSSPRKAVVPARSSLPPNWKQELESLRNGNGSSSSS<br>ASYAAPAAAPARSSSANWRTSPAPS |
| CbbM-EPYC1CIΔCtp | DQSSRYVNLALKEEDLIAGGEHVLCAYIMKPKAGYGYVATAAHFAAESSTGTNVEVCTTDDFTRGVDALVYEVDEARELTKIAYPVALFDRNITDG<br>KAMIASFLTLTMGNNQGMGDVEYAKMHDFYVPEAYRALFDGPSVNISALWKVLGRPEVDGGLVVGTIIKPKLGLRPKPFAEACHAFWLGGDFIK<br>NDEPQGNQPFAPLRDTIALVADAMRRAQDETGEAKLFSANITADDPFEIIARGEYVLETFGENASHVALLVDGYVAGAAAITTARRRFPDNFLHY<br>HRAGHGAVTSPQSKRGYTAFVHCKMARLQGASGIHTGTMGFGKMEGESSDRAIAYMLTQDEAQGPFYRQSWGGMKACTPIISGGMNALRM<br>PGFFENLGNANVILTAGGGAFGHIDGPVAGARSLRQAWQAWRDGVPVLDYAREHKELARAFESFPGDADQIYPGWRKALGVEDTRSALPAAR<br>GSWRDSSSTATVQASRASSATNRASPTRSVLPANWRQELETLRNGNGSSAASSAPAPARSSSASWRDAAPASSAPARSSSASKKAVTPSRSALPS<br>NWKQELESLRSSSPAPASSGSAPAARSSSASWRDAAPASSAPARSSSASKKAVTPSRSALPSNWKQELESLRSSSPAPASSGSAPARSSSASWRDA |

| | |
|---|---|
| | APASSAPARSSSASKKAVTPSRSALPSNWKQELESLRSNSPAPASSGSAPARSSSASWRDAPASSGSAPARSSSASWRDAPASSSSAADKAGTNP<br>WTGKSKPEIKRTPLPADWRKGL |
| CbbM-EPYC1CRΔCtp | DQSSRYVNLALKEEDLIAGGEHVLCAYIMKPKAGYGYVATAAHFAAESSTGTNVEVCTTDDFTRGVDALVYEVDEARELTKIAYPVALFDRNITDG<br>KAMIASFLTLTMGNNQGMGDVEYAKMHDFYVPEAYRALFDGPSVNISALWKVLGRPEVDGGLVVGTIIKPKLGLRPKPFAEACHAFWLGGDFIK<br>NDEPQGNQPFAPLRDTIALVADAMRRAQDETGEAKLFSANITADDPFEIIARGEYVLETFGENASHVALLVDGYVAGAAAITTARRRFPDNFLHY<br>HRAGHGAVTSPQSKRGYTAFVHCKMARLQGASGIHTGTMGFGKMEGESSDRAIAYMLTQDEAQGPFYRQSWGGMKACTPIISGGMNALRM<br>PGFFENLGNANVILTAGGGAFGHIDGPVAGARSLRQAWQAWRDGVPVLDYAREHKELARAFESFPGDADQIYPGWRKALGVEDTRSALPAAR<br>GSWRESSTATVQASRASSATNRVSPTRESVLPANWRQELESLRNGNGSSSAASSAPAPARSSSASWRDAAPASSAPARSSASKKAVTPSRSALPS<br>NWKQELESLRSSSPAPASSAPAPARSSSASQRDAAPASSAPARSSSSKKAVTPSRSALPSNWKQELESLRSSSPAPASSAPAPARSSSASWRDAAPA<br>SSAPARSSSASKKAVTPSRSALPSNWKQELESLRSNSPAPASSAPAPARSSSASWRDAPASSSSSSADKAGTNPWTGKSKPEIKRTALPADWRKGL |
| CbbM-EPYC1EDΔCtp | DQSSRYVNLALKEEDLIAGGEHVLCAYIMKPKAGYGYVATAAHFAAESSTGTNVEVCTTDDFTRGVDALVYEVDEARELTKIAYPVALFDRNITDG<br>KAMIASFLTLTMGNNQGMGDVEYAKMHDFYVPEAYRALFDGPSVNISALWKVLGRPEVDGGLVVGTIIKPKLGLRPKPFAEACHAFWLGGDFIK<br>NDEPQGNQPFAPLRDTIALVADAMRRAQDETGEAKLFSANITADDPFEIIARGEYVLETFGENASHVALLVDGYVAGAAAITTARRRFPDNFLHY<br>HRAGHGAVTSPQSKRGYTAFVHCKMARLQGASGIHTGTMGFGKMEGESSDRAIAYMLTQDEAQGPFYRQSWGGMKACTPIISGGMNALRM<br>PGFFENLGNANVILTAGGGAFGHIDGPVAGARSLRQAWQAWRDGVPVLDYAREHKELARAFESFPGDADQIYPGWRKALGVEDTRSALPAAR<br>GSWRDAPATSAPARPTSAGRASPTRSVLPANWKSELEKLRSSSGNGNGAASAPAPAASARPTSSPKKTATPSRSSLPANWKQELESLRSSSATGSA<br>SSASPSPARSASSSWRDAPSSSSSVAPARSGSAKKPVTPSRSSLPSNWKQELESLRSQSGGSSSSSSAAAAPARSASASWRDAPASSSSSSAAPARS<br>SSAKKPVTPSRSSLPSNWKQELESLRSQSGGSSSSSSAAAAPARSSSASWRDAPASSSSSSSSASKAGSNPWTGKSKPEIKRTTLPADWRKSA |
| CbbM-EPYC1GPΔCtp | DQSSRYVNLALKEEDLIAGGEHVLCAYIMKPKAGYGYVATAAHFAAESSTGTNVEVCTTDDFTRGVDALVYEVDEARELTKIAYPVALFDRNITDG<br>KAMIASFLTLTMGNNQGMGDVEYAKMHDFYVPEAYRALFDGPSVNISALWKVLGRPEVDGGLVVGTIIKPKLGLRPKPFAEACHAFWLGGDFIK<br>NDEPQGNQPFAPLRDTIALVADAMRRAQDETGEAKLFSANITADDPFEIIARGEYVLETFGENASHVALLVDGYVAGAAAITTARRRFPDNFLHY<br>HRAGHGAVTSPQSKRGYTAFVHCKMARLQGASGIHTGTMGFGKMEGESSDRAIAYMLTQDEAQGPFYRQSWGGMKACTPIISGGMNALRM<br>PGFFENLGNANVILTAGGGAFGHIDGPVAGARSLRQAWQAWRDGVPVLDYAREHKELARAFESFPGDADQIYPGWRKALGVEDTRSALPAAR<br>GSWRESSTVTATPAGRSSSAANRVSPTRSVLPANWRQELESLRNGNGNGSSAAAAPAPAPARSASASWRDAPAAAAPARPSSSPKKAVTPSRSS<br>LPANWKQELEALRGGSSSSSASWRTESAPAAAPARSGSKKAVTPSRSSLPANWKQELESMRSASPAPSSAPAAPARSSSASWRSESGSSSSSAAA<br>DKAGTNPWTGKAKVEIKRTALPADWRKGL |
| CbbM-EPYC1TSΔCtp | DQSSRYVNLALKEEDLIAGGEHVLCAYIMKPKAGYGYVATAAHFAAESSTGTNVEVCTTDDFTRGVDALVYEVDEARELTKIAYPVALFDRNITDG<br>KAMIASFLTLTMGNNQGMGDVEYAKMHDFYVPEAYRALFDGPSVNISALWKVLGRPEVDGGLVVGTIIKPKLGLRPKPFAEACHAFWLGGDFIK<br>NDEPQGNQPFAPLRDTIALVADAMRRAQDETGEAKLFSANITADDPFEIIARGEYVLETFGENASHVALLVDGYVAGAAAITTARRRFPDNFLHY<br>HRAGHGAVTSPQSKRGYTAFVHCKMARLQGASGIHTGTMGFGKMEGESSDRAIAYMLTQDEAQGPFYRQSWGGMKACTPIISGGMNALRM<br>PGFFENLGNANVILTAGGGAFGHIDGPVAGARSLRQAWQAWRDGVPVLDYAREHKELARAFESFPGDADQIYPGWRKALGVEDTRSALPAAK<br>GSWRDAPTVTAQPGRAASSAKPTSPTRSVLPANWRQELESLRGGNGNGAAAAPAAAAPRAQSAGWRDAPASAPAASAPMKKTATPARTALP<br>ANWKQELESLRSSSTGGASAAPAAAPARASSASWRDAPAAAPASKSSSPAPAGTNPWTGKSKIEIKRTALPADWRKGL |
| CbbM-EPYC1VAΔCtp | DQSSRYVNLALKEEDLIAGGEHVLCAYIMKPKAGYGYVATAAHFAAESSTGTNVEVCTTDDFTRGVDALVYEVDEARELTKIAYPVALFDRNITDG<br>KAMIASFLTLTMGNNQGMGDVEYAKMHDFYVPEAYRALFDGPSVNISALWKVLGRPEVDGGLVVGTIIKPKLGLRPKPFAEACHAFWLGGDFIK<br>NDEPQGNQPFAPLRDTIALVADAMRRAQDETGEAKLFSANITADDPFEIIARGEYVLETFGENASHVALLVDGYVAGAAAITTARRRFPDNFLHY<br>HRAGHGAVTSPQSKRGYTAFVHCKMARLQGASGIHTGTMGFGKMEGESSDRAIAYMLTQDEAQGPFYRQSWGGMKACTPIISGGMNALRM<br>PGFFENLGNANVILTAGGGAFGHIDGPVAGARSLRQAWQAWRDGVPVLDYAREHKELARAFESFPGDADQIYPGWRKALGVEDTRSALPAAR<br>GSWRESSTVTAQPAGRSSSASARVSPTRSVLPANWRQELESLRNGNGSSSAAAPAPAPAPARSASASWRTEPAPAAASAPSRSPKKAVTPTRSSL<br>PANWKQELESLRGGSSSSSSAPAPAAASAPSRSPKKAVTPTRSSLPANWKQELESLRGGSSSSSSAPAPAAASAPSRSPKKAVTPTRSSLPANWKQ<br>ELESLRGSSSPAPAEAAPARSSSASWRTESPAPAKESSASKAGTNPWTGKSKIEIKRTALPSNWRSGL |
| CbbM-EPYC1VCΔCtp | DQSSRYVNLALKEEDLIAGGEHVLCAYIMKPKAGYGYVATAAHFAAESSTGTNVEVCTTDDFTRGVDALVYEVDEARELTKIAYPVALFDRNITDG<br>KAMIASFLTLTMGNNQGMGDVEYAKMHDFYVPEAYRALFDGPSVNISALWKVLGRPEVDGGLVVGTIIKPKLGLRPKPFAEACHAFWLGGDFIK<br>NDEPQGNQPFAPLRDTIALVADAMRRAQDETGEAKLFSANITADDPFEIIARGEYVLETFGENASHVALLVDGYVAGAAAITTARRRFPDNFLHY<br>HRAGHGAVTSPQSKRGYTAFVHCKMARLQGASGIHTGTMGFGKMEGESSDRAIAYMLTQDEAQGPFYRQSWGGMKACTPIISGGMNALRM<br>PGFFENLGNANVILTAGGGAFGHIDGPVAGARSLRQAWQAWRDGVPVLDYAREHKELARAFESFPGDADQIYPGWRKALGVEDTRSALPAAR<br>GSWRESATVTAQPAGRASSSNRVSPTRSVLPANWRQELESLRNGNGNGAAAAPAPAPAPARSSSASWRSESSAAPAAASTPSRSTKKPVTPTRT<br>SLPANWKQELESLRGSSSSSPAAAAPAPARSSSSPKKAVTPTRSSLPANWKQELESLRGGSSSAASAPAAAAAPAAASAPSRSPKKAVTPTRSSLPA<br>NWKQELESLRGGSSSSSSAPAPAAAPAPARSSSASWRTESPAPANESSSAAAKAGTNPWTGKAKIEIKRTTLPADWRRQL |
| CbbM-EPYC1ΔαΔCtp | DQSSRYVNLALKEEDLIAGGEHVLCAYIMKPKAGYGYVATAAHFAAESSTGTNVEVCTTDDFTRGVDALVYEVDEARELTKIAYPVALFDRNITDG<br>KAMIASFLTLTMGNNQGMGDVEYAKMHDFYVPEAYRALFDGPSVNISALWKVLGRPEVDGGLVVGTIIKPKLGLRPKPFAEACHAFWLGGDFIK<br>NDEPQGNQPFAPLRDTIALVADAMRRAQDETGEAKLFSANITADDPFEIIARGEYVLETFGENASHVALLVDGYVAGAAAITTARRRFPDNFLHY<br>HRAGHGAVTSPQSKRGYTAFVHCKMARLQGASGIHTGTMGFGKMEGESSDRAIAYMLTQDEAQGPFYRQSWGGMKACTPIISGGMNALRM<br>PGFFENLGNANVILTAGGGAFGHIDGPVAGARSLRQAWQAWRDGVPVLDYAREHKELARAFESFPGDADQIYPGWRKALGVEDTRSALPAAR<br>GSWRESSTATVQASRASSATNRVSPTRESVLPANGNGSSSAASSAPAPARSSSASWRDAAPASSAPARSSASKKAVTPSRSALPSNSSPAPASSAP<br>APARSSSASQRDAAPASSAPARSSSSKKAVTPSRSALPSNSSPAPASSAPAPARSSSASWRDAAPASSAPARSSSASKKAVTPSRSALPSNWKQELE<br>SLRSNSPAPASSAPAPARSSSASWRDAPASSSSSSADKAGTNPWTGKSKPEIKRTALPAD |
| CbbM-EPYC1ΔFRΔCtp | DQSSRYVNLALKEEDLIAGGEHVLCAYIMKPKAGYGYVATAAHFAAESSTGTNVEVCTTDDFTRGVDALVYEVDEARELTKIAYPVALFDRNITDG<br>KAMIASFLTLTMGNNQGMGDVEYAKMHDFYVPEAYRALFDGPSVNISALWKVLGRPEVDGGLVVGTIIKPKLGLRPKPFAEACHAFWLGGDFIK<br>NDEPQGNQPFAPLRDTIALVADAMRRAQDETGEAKLFSANITADDPFEIIARGEYVLETFGENASHVALLVDGYVAGAAAITTARRRFPDNFLHY<br>HRAGHGAVTSPQSKRGYTAFVHCKMARLQGASGIHTGTMGFGKMEGESSDRAIAYMLTQDEAQGPFYRQSWGGMKACTPIISGGMNALRM<br>PGFFENLGNANVILTAGGGAFGHIDGPVAGARSLRQAWQAWRDGVPVLDYAREHKELARAFESFPGDADQIYPGWRKALGVEDTRSALPAAR<br>GSWRESSTATVQASRASSAGNGSSSAASSAPAPARSSSASWRDAAPASSAPASSPAPASSAPAPARSSSASQRDAAPASSAPASSPAPASSAPAPA<br>RSSSASWRDAAPASSAPANSPAPASSAPAPARSSSASWRDAPASSSSSSADKAGTNPWTGKSKPEIKRTALPADWRKGL |
| CbbM-EPYC1Anc1ΔCtp | DQSSRYVNLALKEEDLIAGGEHVLCAYIMKPKAGYGYVATAAHFAAESSTGTNVEVCTTDDFTRGVDALVYEVDEARELTKIAYPVALFDRNITDG<br>KAMIASFLTLTMGNNQGMGDVEYAKMHDFYVPEAYRALFDGPSVNISALWKVLGRPEVDGGLVVGTIIKPKLGLRPKPFAEACHAFWLGGDFIK<br>NDEPQGNQPFAPLRDTIALVADAMRRAQDETGEAKLFSANITADDPFEIIARGEYVLETFGENASHVALLVDGYVAGAAAITTARRRFPDNFLHY<br>HRAGHGAVTSPQSKRGYTAFVHCKMARLQGASGIHTGTMGFGKMEGESSDRAIAYMLTQDEAQGPFYRQSWGGMKACTPIISGGMNALRM<br>PGFFENLGNANVILTAGGGAFGHIDGPVAGARSLRQAWQAWRDGVPVLDYAREHKELARAFESFPGDADQIYPGWRKALGVEDTRSALPAAR<br>GSWRESSTATVQASRSASAANRVSPTRSVLPANWRQELESLRNGNGSSSSASSAPAPARSSSASWRDAPASSAPARSSSASKKAVTPSRSSLPAN<br>WKQELESLRSSSPAPASSSAAPARSSSASWRDAPASSAPARSSSASKKPVTPSRSSLPSNWKQELESLRSSSPAPASSSAAPARSSSASWRDAPASS<br>APARSSSASKKAVTPSRSSLPSNWKQELESLRSSSPAPASSSAAPARSSSASWRDAPASSSSSSADKAGTNPWTGKSKVEIKRTTLP |
| CbbM-EPYC1Anc2ΔCtp | DQSSRYVNLALKEEDLIAGGEHVLCAYIMKPKAGYGYVATAAHFAAESSTGTNVEVCTTDDFTRGVDALVYEVDEARELTKIAYPVALFDRNITDG<br>KAMIASFLTLTMGNNQGMGDVEYAKMHDFYVPEAYRALFDGPSVNISALWKVLGRPEVDGGLVVGTIIKPKLGLRPKPFAEACHAFWLGGDFIK<br>NDEPQGNQPFAPLRDTIALVADAMRRAQDETGEAKLFSANITADDPFEIIARGEYVLETFGENASHVALLVDGYVAGAAAITTARRRFPDNFLHY<br>HRAGHGAVTSPQSKRGYTAFVHCKMARLQGASGIHTGTMGFGKMEGESSDRAIAYMLTQDEAQGPFYRQSWGGMKACTPIISGGMNALRM<br>PGFFENLGNANVILTAGGGAFGHIDGPVAGARSLRQAWQAWRDGVPVLDYAREHKELARAFESFPGDADQIYPGWRKALGVEDTRSALPAAR<br>GSWRDSSTVTATQPGRSSSAANRVSPTRSVLPANWRQELESLRNGNGNGSSAASAPAPARSSSASWRDAPAPAAPARPSSSPKKTVTPSRSSLP |

| | |
|---|---|
| | ANWKQELESLRSSSSGSASSSAPSARSSSSSWRDPPASSAPARSRSASKKPVTPSRSSLPSNWKQELESLRGGSPAPSSSAAAPARSASASWRDPA<br>ASSAPARSSSASKKAVTPSRSSLPSNWKQELESLRSSSPAPASAAAAPARSSSASWRDAPASSSSSSADKAGTNPWTGKSKIEIKRTTLP |
| CbbM-EPYC1Anc3ΔCtp | DQSSRYVNLALKEEDLIAGGEHVLCAYIMKPKAGYGYVATAAHFAAESSTGTNVEVCTTDDFTRGVDALVYEVDEARELTKIAYPVALFDRNITDG<br>KAMIASFLTLTMGNNQGMGDVEYAKMHDFYVPEAYRALFDGPSVNISALWKVLGRPEVDGGLVVGTIIKPKLGLRPKPFAEACHAFWLGGDFIK<br>NDEPQGNQPFAPLRDTIALVADAMRRAQDETGEAKLFSANITADDPFEIIARGEYVLETFGENASHVALLVDGYVAGAAAITTARRRFPDNFLHY<br>HRAGHGAVTSPQSKRGYTAFVHCKMARLQGASGIHTGTMGFGKMEGESSDRAIAYMLTQDEAQGPFYRQSWGGMKACTPIISGGMNALRM<br>PGFFENLGNANVILTAGGGAFGHIDGPVAGARSLRQAWQAWRDGVPVLDYAREHKELARAFESFPGDADQIYPGWRKALGVEDTRSALPAAR<br>GSWRESSTATVQASRASSATNRVSPTRSVLPANWRQELESLRNGNGSSSAASSAPAPARSSSASWRDAAPASSAPARSSSASKKAVTPSRSALPS<br>NWKQELESLRSSSPAPASSASAPARSSSASWRDAAPASSAPARSSSASKKAVTPSRSALPSNWKQELESLRSSSPAPASSASAPARSSSASWRDAA<br>PASSAPARSSSASKKAVTPSRSALPSNWKQELESLRSNSPAPASSASAPARSSSASWRDAPASSSSSSADKAGTNPWTGKSKPEIKRTALP |
| CbbM-EPYC1Anc4ΔCtp | DQSSRYVNLALKEEDLIAGGEHVLCAYIMKPKAGYGYVATAAHFAAESSTGTNVEVCTTDDFTRGVDALVYEVDEARELTKIAYPVALFDRNITDG<br>KAMIASFLTLTMGNNQGMGDVEYAKMHDFYVPEAYRALFDGPSVNISALWKVLGRPEVDGGLVVGTIIKPKLGLRPKPFAEACHAFWLGGDFIK<br>NDEPQGNQPFAPLRDTIALVADAMRRAQDETGEAKLFSANITADDPFEIIARGEYVLETFGENASHVALLVDGYVAGAAAITTARRRFPDNFLHY<br>HRAGHGAVTSPQSKRGYTAFVHCKMARLQGASGIHTGTMGFGKMEGESSDRAIAYMLTQDEAQGPFYRQSWGGMKACTPIISGGMNALRM<br>PGFFENLGNANVILTAGGGAFGHIDGPVAGARSLRQAWQAWRDGVPVLDYAREHKELARAFESFPGDADQIYPGWRKALGVEDTRSALPAAR<br>GSWRDAPTVTAQPGRSSSAANRVSPTRSVLPANWRQELESLRNGNGNGSSAASAPAPARSSSASWRDAPAAPARPSSSPKKTATPSRSSLPAN<br>WKQELESLRSSSTGSASSAAPSARSSSSSWRDPSSSSSVAPARSGSKKPVTPSRSSLPSNWKQELESLRGGSSSSSSAAAAPARSASASWRDPAAS<br>SAPARSSSSKKAVTPSRSSLPSNWKQELESLRSSSPAPASAAAAPARSSSASWRDAPASSSSSSASKAGTNPWTGKSKIEIKRTTLP |
| CbbM-EPYC1Anc5ΔCtp | DQSSRYVNLALKEEDLIAGGEHVLCAYIMKPKAGYGYVATAAHFAAESSTGTNVEVCTTDDFTRGVDALVYEVDEARELTKIAYPVALFDRNITDG<br>KAMIASFLTLTMGNNQGMGDVEYAKMHDFYVPEAYRALFDGPSVNISALWKVLGRPEVDGGLVVGTIIKPKLGLRPKPFAEACHAFWLGGDFIK<br>NDEPQGNQPFAPLRDTIALVADAMRRAQDETGEAKLFSANITADDPFEIIARGEYVLETFGENASHVALLVDGYVAGAAAITTARRRFPDNFLHY<br>HRAGHGAVTSPQSKRGYTAFVHCKMARLQGASGIHTGTMGFGKMEGESSDRAIAYMLTQDEAQGPFYRQSWGGMKACTPIISGGMNALRM<br>PGFFENLGNANVILTAGGGAFGHIDGPVAGARSLRQAWQAWRDGVPVLDYAREHKELARAFESFPGDADQIYPGWRKALGVEDTRSALPAAR<br>GSWRDSSTVTATPAGRSSSAANRVSPTRSVLPANWRQELESLRNGNGNGSSAASAPAPARSSSASWRDAPAPAAPARPSSSPKKAVTPSRSSLPA<br>NWKQELESLRSGSSGSSSSSAPAARSSSSKKPVTPSRSSLPANWKQELESLRGGSSSSASAPARSASASWRDPAAASAPARSSSGSKKAVTPSRSSL<br>PANWKQELESLRSSSPAPASAAAAPARSSSASWRDSPASSSSSSADKAGTNPWTGKSKIEIKRTTLP |
| CbbM-EPYC1Anc6ΔCtp | DQSSRYVNLALKEEDLIAGGEHVLCAYIMKPKAGYGYVATAAHFAAESSTGTNVEVCTTDDFTRGVDALVYEVDEARELTKIAYPVALFDRNITDG<br>KAMIASFLTLTMGNNQGMGDVEYAKMHDFYVPEAYRALFDGPSVNISALWKVLGRPEVDGGLVVGTIIKPKLGLRPKPFAEACHAFWLGGDFIK<br>NDEPQGNQPFAPLRDTIALVADAMRRAQDETGEAKLFSANITADDPFEIIARGEYVLETFGENASHVALLVDGYVAGAAAITTARRRFPDNFLHY<br>HRAGHGAVTSPQSKRGYTAFVHCKMARLQGASGIHTGTMGFGKMEGESSDRAIAYMLTQDEAQGPFYRQSWGGMKACTPIISGGMNALRM<br>PGFFENLGNANVILTAGGGAFGHIDGPVAGARSLRQAWQAWRDGVPVLDYAREHKELARAFESFPGDADQIYPGWRKALGVEDTRSALPAAR<br>GSWRESSTVTATPAGRSSSAANRVSPTRSVLPANWRQELESLRNGNGNGSSAASAPAPAPARSASASWRDAPAPAAPARPSSSPKKAVTPSRSSL<br>PANWKQELESLRGGSSSSSSSAAPAARSSSSKKPVTPTRSSLPANWKQELESLRGGSSSSASAPAPAAASAPARSGSKKAVTPSRSSLPANWKQEL<br>ESLRSSSPAPASAPAAPARSSSASWRSESPASSSSSSADKAGTNPWTGKAKIEIKRTTLP |
| CbbM-EPYC1Anc7ΔCtp | DQSSRYVNLALKEEDLIAGGEHVLCAYIMKPKAGYGYVATAAHFAAESSTGTNVEVCTTDDFTRGVDALVYEVDEARELTKIAYPVALFDRNITDG<br>KAMIASFLTLTMGNNQGMGDVEYAKMHDFYVPEAYRALFDGPSVNISALWKVLGRPEVDGGLVVGTIIKPKLGLRPKPFAEACHAFWLGGDFIK<br>NDEPQGNQPFAPLRDTIALVADAMRRAQDETGEAKLFSANITADDPFEIIARGEYVLETFGENASHVALLVDGYVAGAAAITTARRRFPDNFLHY<br>HRAGHGAVTSPQSKRGYTAFVHCKMARLQGASGIHTGTMGFGKMEGESSDRAIAYMLTQDEAQGPFYRQSWGGMKACTPIISGGMNALRM<br>PGFFENLGNANVILTAGGGAFGHIDGPVAGARSLRQAWQAWRDGVPVLDYAREHKELARAFESFPGDADQIYPGWRKALGVEDTRSALPAAR<br>GSWRESSTVTAQPAGRSSSASNRVSPTRSVLPANWRQELESLRNGNGNGSAAAPAPAPAPARSSSASWRESPAPAAASTPSRSPKKAVTPTRSSL<br>PANWKQELESLRGGSSSSSSAAAPAPARSSSSKKAVTPTRSSLPANWKQELESLRGGSSSSASAPAPAAASAPSRSPKKAVTPTRSSLPANWKQEL<br>ESLRGSSSPAPASAPAPARSSSASWRTESPAPANESSASKAGTNPWTGKAKIEIKRTTLP |

**Supplementary table 2 Parameter estimates from mass action-mass transfer model fit to observed data.** Models were fitted to 0.3 mM and 1 mM RuBP aerobic time curve data. For both situations (varying partitioning or $K_M$), substrate concentrations, active site content and phase volume were varied as well to attain good model fit. For $K_M$ variation only the $K_M$s inside the condensates were considered. Enzyme partitioning was assumed at 99%, condensate radius at average 10 µm, diffusion coefficient of substrates at 1e-11 µm²/s, as measured for other small molecules previously[41].

| | Mass action-mass transfer partitioning | | | Mass action-mass transfer catalytic parameters inside the condensates | | |
|---|---|---|---|---|---|---|
| | $K(CO_2)$ | $K(O_2)$ | K(RuBP) | $K_M(CO_2)$ | $K_M(O_2)$ | $K_M$(RuBP) |
| 0.3 mM RuBP | 171 | 47 | 1 | 18 | 4032 | 781 |
| 1 mM RuBP | 96 | 11 | 0.4 | 66 | 768 | 806 |

**Supplementary table 3 Percentage Identity of ancestors inferred via mtInv versus JJT.** The percentage identity of sequences inferred with two different models to test for the ASR robustness is shown.

| Protein | Percentage Identity [%] |
|---|---|
| Anc1 | 100 |
| Anc2 | 99.03 |
| Anc3 | 99.04 |
| Anc4 | 99.34 |
| Anc5 | 97.91 |
| Anc6 | 98.56 |
| Anc7 | 98.56 |

**Supplementary table 4 Percentage Identity of natural variants and ancestors with respect to the EPYC1 from *C. reinhardtii*.**

| Protein | Percentage Identity [%] |
|---|---|
| *Chlamydomonas reinhardtii* | 100 |
| *Chlamydomonas schloesseri* | 82.818 |
| *Chlamydomonas incerta* | 83.013 |
| *Volvox africanus* | 57.823 |
| *Volvox carteri* | 55.96 |
| *Edaphochlamys debaryana* | 55.049 |
| *Gonium pectoral* | 56.277 |
| *Astrephomene gubernaculifera* | 58.824 |
| *Tetrabaena socialis* | 56.977 |
| Anc1 | 84.099 |
| Anc2 | 80.702 |
| Anc3 | 97.544 |
| Anc4 | 76.491 |
| Anc5 | 68.31 |
| Anc6 | 58.188 |
| Anc7 | 57.377 |

**Supplementary table 5 X-ray data collection and structure refinement statistics.**

| Sample | Inactive form of CbbM | Activated form of CbbM |
|---|---|---|
| Accession | PDB 8S20 | PDB 8S21 |
| Ligands | - | $Mg^{2+}$, CABP |
| **Data collection** | | |
| Beamline | PETRA III - P13 | PETRA III - P13 |
| Wavelength (Å) | 0.9762 | 0.9762 |
| Space Group | $P\ 4_1\ 2_1\ 2$ | $P\ 3_2\ 2\ 1$ |
| Unit cell dimensions | | |
| a, b, c (Å) | 78.95, 78.95, 277.20 | 133.84, 133.84, 151.77 |
| α, β, γ (°) | 90.00, 90.00, 90.00 | 90.00, 90.00, 90.00 |
| Resolution (Å) | 29.78 - 2.50 (2.56 - 2.50) | 29.60 - 2.40 (2.46 - 2.40) |
| Unique reflections | 31424 (2290) | 61779 (4526) |
| Multiplicity | 13.26 (13.85) | 20.36 (20.91) |
| Completeness (%) | 99.9 (100.0) | 99.9 (100.0) |
| $I / \sigma I$ | 22.7 (2.8) | 12.6 (2.3) |
| $R_{meas}$ | 0.116 (1.366) | 0.292 (2.392) |
| $CC_{1/2}$ | 0.999 (0.780) | 0.997 (0.737) |
| **Refinement** | | |
| $R_{work}$ / $R_{free}$ | 0.208 / 0.239 | 0.2024 / 0.2292 |
| RMS bonds | 0.002 | 0.002 |
| RMS angles | 0.445 | 0.514 |
| Ramachandran | | |
| favored (%) | 97.40 | 97.47 |
| allowed (%) | 2.23 | 2.53 |
| outliers (%) | 0.37 | 0.00 |
| Rotamer outliers (%) | 0.16 | 0.29 |
| Number of atoms | 6471 | 7482 |
| Protein | 6337 | 7000 |
| Ligands | - | 438 |
| Solvent | 134 | 44 |
| Average B-factor | 59.31 | 41.49 |
| Protein | 59.42 | 41.38 |
| Ligands | - | 36.85 |
| Solvent | 53.78 | 43.64 |

**Supplementary table 6 CryoEM data collection, refinement, and model statistics.**

| **Data collection** | |
|---|---|
| Model | CbbM-EPYC1$_{CR}$Δα |
| Accessions | PDB 8S22, EMD-19652 |
| Microscope | Titan Krios G3i |
| Voltage (kV) | 300 |
| Camera | K3 |
| Magnification | 130,000 |
| Pixel size at detector (Å/pixel) | 0.655 |
| Total electron exposure ($e^-$/Å$^2$) | 60 |
| Frames per exposure | 60 |
| Defocus range (µm) | -1.2 to -2.4 |
| Automation software | EPU, CryoSPARC |
| Micrographs collected (no.) | 11525 |
| Micrographs used (no.) | 7116 |
| Total extracted particles (no.) | 45233 |
| Final particles (no.) | 18706 |
| Point-group | *C* 2 |
| Resolution (global, Å) | 3.4 |
| FSC$_{0.143}$ | 4.3 / 3.4 (unmasked / masked) |
| Resolution range (local, Å) | 2.2 – 12.0 (according to local resolution histogram) |
| Map sharpening B-factor (Å$^2$) | -98.7 |
| Map sharpening methods | CryoSPARC sharpening |
| **Model Refinement** | |
| Refinement package | Phenix |
| method | real space |
| resolution cutoff | 3.38 |
| Protein residues | 910 (2×modified lysine - KCX) |
| Ligands | 2× $Mg^{2+}$, 2× CABP |
| Model-Map scores | |
| CC$_{Volume}$ | 0.82 |
| avg. FSC$_{0.143}$ | 3.43 / 3.38 (unmasked / masked) |
| B-factors (Å$^2$) (min / max / mean) | |
| Protein residues | 55.37 / 162.98 / 94.25 |
| Ligands | 90.19 / 138.27 / 111.78 |
| R.M.S. Bond lengths (Å) | 0.005 |
| R.M.S. Bond angles (°) | 0.665 |
| **Validation** | |
| MolProbity score | 1.72 |
| CaBLAM outliers | 1.44 |
| Clashscore | 9.11 |
| Poor rotamers (%) | 0.73 |
| C-beta deviations | 0.00 |
| Ramachandran favored (%) | 96.79 |
| Ramachandran outliers (%) | 0.00 |
